# Varying parameter ranges alters both Partial Rank Correlation Coefficient results and phenomenological behavior when modeling the epithelial mesenchymal transition

**DOI:** 10.64898/2026.06.05.730399

**Authors:** Kelsey I. Gasior

**Author notes:** Corresponding authors *Email address:* (Kelsey I. Gasior).

## Abstract

Partial Rank Correlation Coefficient (PRCC), usually performed following Latin Hyper-cube Sampling (LHS), is a global sensitivity analysis that quantifies the monotonic relationship between model parameters and the desired output. To carry out this analysis, a range of acceptable parameter values must be known or estimated. However, within a biological context, approximating these values may be difficult. Parameter values and ranges can be taken from different organisms or systems or be estimated to produce qualitative phenomena in the model. Using a mathematical model of the epithelial mesenchymal transition (EMT) as a test case, this work examines how the parameter ranges chosen prior to analysis can influence LHS-PRCC results and shape subsequent analysis interpretations. Previous LHS-PRCC analysis of this model restricted parameters to ±10% of their original value, which limits the scope and interpretability of parameter influence. Such a small range assumes, in the biological sense, that parameters are well-measured with little variability. Here, this work extends the previous analysis and explores several parameter ranges (±25%, ±50% of the original value). This work also tests whether, within the ±10%, ±25% and ±50% parameter ranges, the bistable switch present in the original model are maintained. Ultimately, this work showcases how a choice made prior to analysis, such as the accepted parameter ranges for biological rates and values in complex dynamical systems can influence sensitivity analysis results and interpretability. Additionally, these choices can have hidden consequences, such as the loss of phenomenological behavior. Thus, explicit prior knowledge about the appropriate parameter values is needed before using analysis to guide future experiments and model development.

## 2. Introduction

Carcinomas, derived from epithelial cells, are the most common type of tumor [1; 2]. Epithelial cells are defined by their highly adhesive nature and this adhesion is due to the transmembrane protein, E-cadherin [2; 3]. Membrane-bound E-cadherin interacts with other E-cadherin molecules on neighboring cells while simultaneously being anchored to the cytoskeleton on the intracellular side of the membrane via its bonds with members of the catenin family, such as *α*-, *β*-, and *γ*-catenin [2]. However, for metastasis formation, cells must leave the tumor microenvironment and migrate elsewhere in the body, requiring the cells to lose their adhesion [4]. This happens through a process called the epithelial mesenchymal transition (EMT). During EMT, cells lose their adhesive properties and acquire the invasive and migratory properties associated with mesenchymal cells [3; 5]. There are many pathways that can activate EMT and it has been hypothesized that there is an underlying bistable switch with respect to these different exogenous activation mechanisms [6–9]. By undergoing the switch, the cell can migrate away from the tumor and maintain the mesenchymal phenotype, even after losing the original activating signal.

EMT, associated intracellular components, and the underlying switch have been extensively studied in the literature [7; 10–13]. One recent study examined EMT experimentally in two different human cell lines: colon carcinoma cells (SW480) and breast carcinoma cells (MCF7). The authors studied how TGF-*β* and cellular contact impact the activation of EMT in both cell lines. Exogenous TGF-*β* is a known activator of EMT in carcinoma cells and cellular contact has been shown to encourage the epithelial phenotype. Through their experimental research, the authors showed how the application of TGF-*β* to cell cultures at different confluences could encourage EMT in both cell lines. Additionally, the application of TGF-*β* to the MCF7 cells produced bistable behavior, as indicated by the EMT-associated changes in both the expression of E-cadherin and Slug, as well as the phenotype present. It was also noted that MCF7 cells without any cellular contact and in the absence of exogenous TGF-*β* also exhibited the mesenchymal phenotype, indicating that EMT and the switch could occur with respect to cellular contact [14].

The authors then constructed a mathematical model of EMT in MCF7 breast carcinoma cells to understand the link between changes to intracellular components and behavioral changes. This work included a two-equation ordinary differential equation (ODE) model of E-cadherin, a classic marker associated with the epithelial state, and Slug, a transcription factor downstream of TGF-*β*. Based on the phenomenological behavior observed experimentally, the authors proposed the existence of two bistable switches in MCF7 cells: an irreversible switch with respect to TGF-*β* and a reversible switch due to a loss of cellular contact. This model showed that cells without any contact began as mesenchymal cells and remained mesenchymal cells in the presence of exogenous TGF-*β*. Cells with little-to-moderate levels of contact (1 or 2 neighbors) could undergo the irreversible bistable switch with respect to TGF-*β* but, cells with high levels of contact, such as 6 neighboring cells, were anchored in place by their neighbors and could not undergo the switch with respect to TGF-*β*. With this model, the authors found that a cell with 6 neighbors could undergo a TGF-*β*-induced switch was if that cell simultaneously lost cellular contact as it was exposed to the exogenous factor. For these high-contact cells, both factors must work together to activate EMT. Thus, this model showed that tumor cells most likely to undergo EMT due to exogenous TGF-*β* exposure are those with little cellular contact. Without the loss of these low-contact cells, cells towards the center of the tumor cannot transition, indicating that metastasis prevention should focus on maintaining the epithelial phenotype in tumor edge cells. Next, given that the parameters were estimated to capture the bistable, phenomenological behavior present in the MCF7 data, the authors then nondimensionalized the model, analyzed it, and compared their results to the low-confluence MCF7 experimental results. This comparison found that the changes to E-cadherin and Slug in the model were within the same range of E-cadherin and Slug mRNA changes found experimentally [8].

Phenomenological models, such as this one crafted around MCF7 data, require methodical analysis to understand how model parameters based on rates and concentrations are linked to observed changes, such as intracellular component concentrations and phenotypic shifts. Global sensitivity analysis (GSA) is a powerful tool for understanding how uncertainty within a model is a result of the parameters and quantifying the impact of these parameters on the desired outputs [15; 16]. Parameter values maybe uncertain due to a lack of data or gaps in existing data (epistemic) or inherent variations in the modeling structure and GSA can determine the impact of one parameter or a group of parameters on outputs. Unlike local sensitivity analysis, which requires parameters be varied one-at-a-time, GSA allows for model parameters to be varied simultaneously [17; 18]. However, GSA methods require more input samples, which can become an issue if each input significantly increases the computational cost [17].

One popular type of global sensitivity analysis that can be used is Partial Rank Correlation Coefficient (PRCC). Global sensitivity analyses, such as Sobol’ analysis, can be computationally expensive. Comparatively PRCC has a moderate computational cost [19]. Further, it has been shown that there is a correlation between the results of PRCC and total Sobol’, making it an accessible alternative for analysis [17; 20]. PRCC is carried out following a sampling methodology, such as Latin Hypercube Sampling (LHS) [19; 21; 22]. LHS is a type of Monte Carlo Sampling where each parameter range is sampled *N* times [23]. A parameter value is used once and the values are assembled into *N* parameter vectors to produce the necessary outputs [23; 24]. PRCC can then rank the input values for each parameter and the model outputs and quantify the monotonic relationship between them via the correlation coefficient (*ρ*) [19; 25]. Additionally, PRCC can provide directionality to this relationship (− 1 ≤ *ρ* ≤ 1). Throughout the literature, a value of |*ρ*| *>* 0.5 has been used as a cutoff for sensitivity [7–9; 19; 26–29].

Following the establishment of the model in [8], extensive analysis via LHS-PRCC was performed on the model to determine how both modeling choices and parameter values impacted measurable, experimental quantities [9]. Given that the original model was nondimensionalized, subsequent analysis worked to determine the impact of that nondimensionalization on the results of PRCC [8; 9]. To mimic the heterogeneity of the tumor microenvironment and the exposure to the EMT-inducing factor, TGF-*β*, this analysis was carried out over four different levels of cellular contact and two different levels of TGF-*β*. Using a parameter range of ±10% of the given model value for construction of the Latin Hypercube, this analysis compared the PRCC results from 7 different iterations of nondimensionalization to each other and analysis of the original, dimensional model under 8 different exogenous conditions. The ±10% parameter range allowed LHS-PRCC analysis to determine how small changes in the original bifurcation-producing values might impact the PRCC results for both the dimensional and nondimensional models. Ultimately, the results of these works showed that differences in PRCC results can stem from biological conditions inherent to the system, such as exogenous factors activating EMT, and simplifications introduced by the analyst, such as nondimensionalization. Under different EMT-activating conditions, PRCC highlighted different parameters affecting the long-term values of intracellular EMT-associated components. But these relationships are altered or lost upon nondimensionalization which can exclude important parameters from analysis altogether.

While this analysis offered new insights on the impact of nondimensionalization, the choice of a small parameter range (±10%) around the original model value also imposed limitations on the understanding of the model from PRCC [9]. A small parameter range around the given model value indicates a known or measured quantity with little deviation or error in measurements. But, depending on the source of the parameter values, these ranges can be much larger or possibly unknown. Parameter values chosen from the literature can, potentially, be from different cell cultures, organism, or model organisms. Additional variation can occur depending on the experimental treatment conditions and potential caveats present during data collection can alter the parameter range. Finally, some parameter values are chosen to help model phenomenological behavior. Any combination of these possibilities could dictate the use of a parameter range that is inappropriately sized or, worse, holding a parameter constant under the incorrect assumption that it is known. Previous work on sensitivity analysis has shown that the size of the parameter range can affect the results [30], but it can still be difficult to fully grasp how much influence the parameter choice can have in complicated, multi-scale systems that are subjected to a variety of external conditions. It can be additionally frustrating to understand what the deviations in PRCC results can mean when attempting to use the results to guide additional experiments as part of a computational-experimental feedback loop.

This work seeks to show the issues that can arise from unknown parameter ranges when analyzing biomathematical models. Using the original EMT model established in [8], the analysis performed in [9] chose a range of ±10% around the original bifurcation-producing values, suggesting a well-measured and known parameter value. However, these values were chosen to capture the phenomenological EMT behavior observed in MCF7 cells, rather than exactly measured [8; 14]. Given the robust range of each parameter that could produce this behavior, the ±10% range is too narrow. As a first step, this work extends the PRCC analysis to include larger parameter ranges (±25%, ±50%) around the original model value and carries out analysis at two different levels of exogenous TGF-*β* and four levels of cell-cell contact. The same quantity of interest (QOI) is used for analysis: the steady state values of the E-cadherin protein and the transcription factor Slug. These values were chosen as they could be measured experimentally, making this analysis applicable and helpful when suggesting new experimental directions based on modeling results. Additionally, this work examines how analytical choices can affect phenomenological behavior. The original model sought to capture the bistability of EMT in MCF7 cells [8; 14] but deviating from these values simultaneously, as occurs when building a Latin Hypercube, may result in a loss of the bifurcation. The robustness of this phenomenological behavior is during analysis is impacted by the size of the parameter range chosen. Further, failure to consider how this behavior changes can impact the meaning and interpretability of PRCC results.

Collectively, the results here show that the parameter range is important when using LHS-PRCC analysis to interpret the impact of specific model parameters on measurable experimental values but, even with careful analysis, hidden consequences can occur. Different parameter ranges for LHS-PRCC can result in shifts or loss in sensitivity for certain parameters within the tumor and the original bifurcation in the model is not preserved when analysis is carried out. Through this analysis there are similar patterns that emerge for the parameters influencing the steady state levels of E-cadherin and Slug, as well as the loss of bistability, across all three ranges (±10%, ±25%, ±50%). But these similarities are only visible because analysis is conducted as part of a larger attempt to understand hidden underlying effects. Performing only one analysis with set ranges could lead to an incorrect conclusion on what parameters intracellular EMT-associated components and these incorrect results could impact biological experiments performed on the recommendations of the analysis. Ultimately, this work shows how user choices, such as parameter range, compounded with complex multi-scale dynamics in biological systems, can have consequences for the interpretation of the analysis.

## 3. Methods

### 3.1. Mathematical Model of E-cadherin and Slug

Gasior et al. put forth a two-equation single-cell model of EMT in the breast carcinoma cell line, MCF7. This model focused on a classic marker of the epithelial phenotype, the transmembrane protein E-cadherin, and a transcription factor associated with the mesenchymal phenotype, Slug [8]. A schematic of this model is shown in Figure 1, the ordinary differential equation model for E-cadherin (*E*) and Slug (*S*) is shown in Equations 1-2, and parameter values are found in Table 1. In addition to production (*α*_*i*_, *i* = 1, 2) and degradation (*β*_*i*_, *i* = 1, 2), this model examines the changes in E-cadherin and Slug in response to changes in two exogenous factors capable of activating EMT: TGF-*β* (*T*) and cellular contact (*C*).

**Table 1:** Parameter values for model in Equations 1-2 and originally proposed by [8].

| Parameter | Definition | Assumed Value | Units | Source |
| --- | --- | --- | --- | --- |
| $\alpha_1$ | E-cadherin production | 0.19355 | $\frac{\text{ng}}{\text{mL}\cdot\text{min}}$ | Estimated |
| $\alpha_2$ | Slug production | 0.080 | $\frac{\text{ng}}{\text{mL}\cdot\text{min}}$ | Estimated |
| $\beta_1$ | E-cadherin degradation | 6.035 | $\frac{1}{\text{min}}$ | Estimated |
| $\beta_2$ | Slug degradation | 0.830 | $\frac{1}{\text{min}}$ | Estimated |
| $k_0$ | Rate at which E-cadherin moves to the membrane due to cell-cell contact | 0.1245 | $\frac{\text{ng}}{\text{mL}\cdot\text{min}}$ | Estimated |
| $k_1$ | Rate of Slug suppression from membrane-bound E-cadherin activation | 0.080 | $\frac{\text{ng}}{\text{mL}\cdot\text{min}}$ | Estimated |
| $k_2$ | Rate of Slug upregulation via TGF-β | 0.040 | $\frac{\text{ng}}{\text{mL}\cdot\text{min}}$ | Estimated |
| $IC_S$ | Half maximal concentration of Slug required to inhibit E-cadherin production | 0.01920 | $\frac{\text{ng}}{\text{mL}}$ | Estimated |
| $IC_E$ | Half maximal concentration of E-cadherin required to inhibit Slug upregulation | 0.010 | $\frac{\text{ng}}{\text{mL}}$ | Estimated |
| $IC_T$ | Half maximal concentration of TGF-β required to activate Slug | $3.64 \times 10^{-6}$ | $\frac{\text{ng}}{\text{mL}}$ | Estimated |
| $IC_C$ | Half maximal number of cells required to draw E-cadherin to the membrane for activation | 2.17 | cells | Estimated |
| $n_1$ | Hill Coefficient | 3 | — | Estimated |
| $n_2$ | Hill Coefficient | 4 | — | Estimated |
| $n_3$ | Hill Coefficient | 2 | — | Estimated |
| $n_4$ | Hill Coefficient | 3 | — | Estimated |

**Figure 1:**
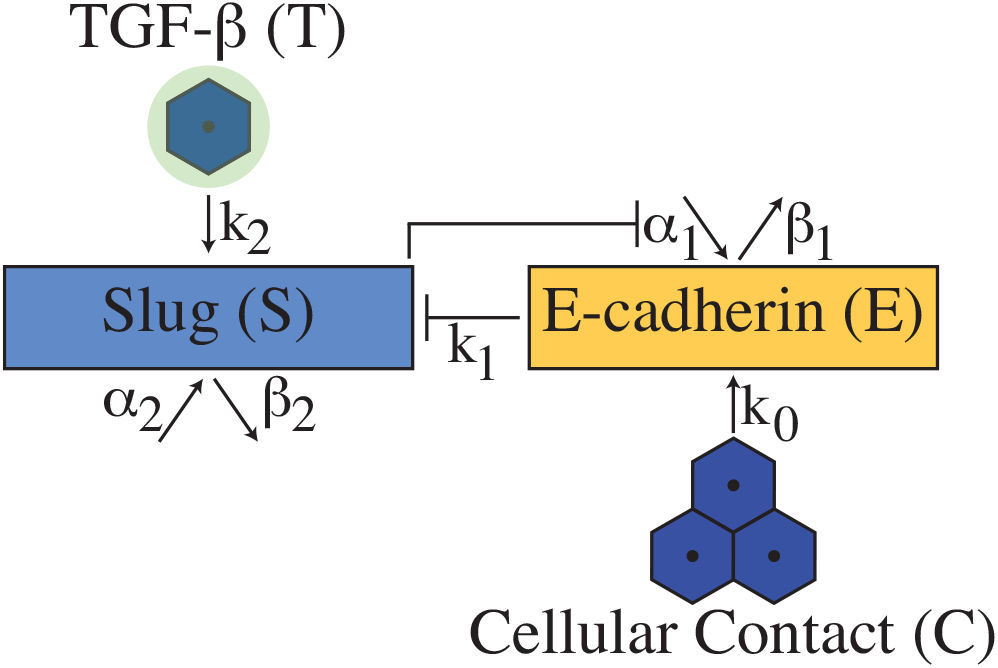
In the presence of neighboring epithelial cells (*C >* 0), E-cadherin translocates to the membrane to form the intercellular interactions with E-cadherin on neighboring cells responsible for cellular adhesion. E-cadherin simultaneously forms intracellular bonds with *β*-catenin. This interaction prevents *β*-catenin from translocating to the nucleus, suppressing Slug activation. The addition of exogenous TGF-*β* (*T*) activates an intracellular signaling cascade, upregulating Slug, which then suppresses E-cadherin production. Loss of E-cadherin allows for *β*-catenin accumulation and translocation to the nucleus, further activating Slug [8; 9]

Without cellular contact, E-cadherin is capable of being endocytosed into the cell. In the presence of cellular contact (*C*), E-cadherin moves to the membrane to form intercellular bonds with neighboring epithelial cells 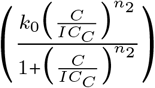 [2; 31]. In this model, *C* is a continuous input variable, meaning it is possible to have fractional levels of contact. To anchor it to the membrane, E-cadherin binds with members of the catenin family, such as *β*-catenin, on the intracellular side of the membrane, linking it with the actin cytoskeleton [2]. This interaction with E-cadherin prevents *β*-catenin from translocating to the nucleus to activate Slug. The sequestration of *β*-catenin at the membrane that suppresses Slug activation is modeled by 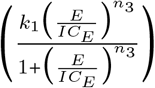 in Equation 2 [31–33].

If TGF-*β* is released from the surrounding microenvironment, molecules can to bind with the receptors on the cellular membrane and activate intracellular signaling cascades that upregulate the transcription factor Slug 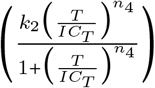 [34–36]. Slug then inhibits the transcription of E-cadherin 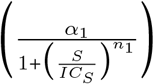, reducing the pool of E-cadherin available for eventual intercellular interactions [33; 37]. With the upregulation of Slug and the loss of E-cadherin available for cell-cell contact, the cell can undergo EMT and acquire the invasive and migratory properties associated with mesenchymal cells.

For all subsequent analysis presented here, four levels of cellular contact (*C* = 0, 1, 2, 6 cells) and two levels of exogenous TGF-*β* 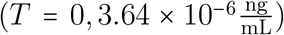 were considered and are shown in Figure 2.

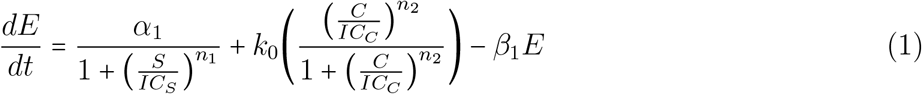

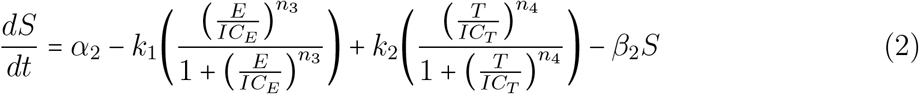

**Figure 2:**
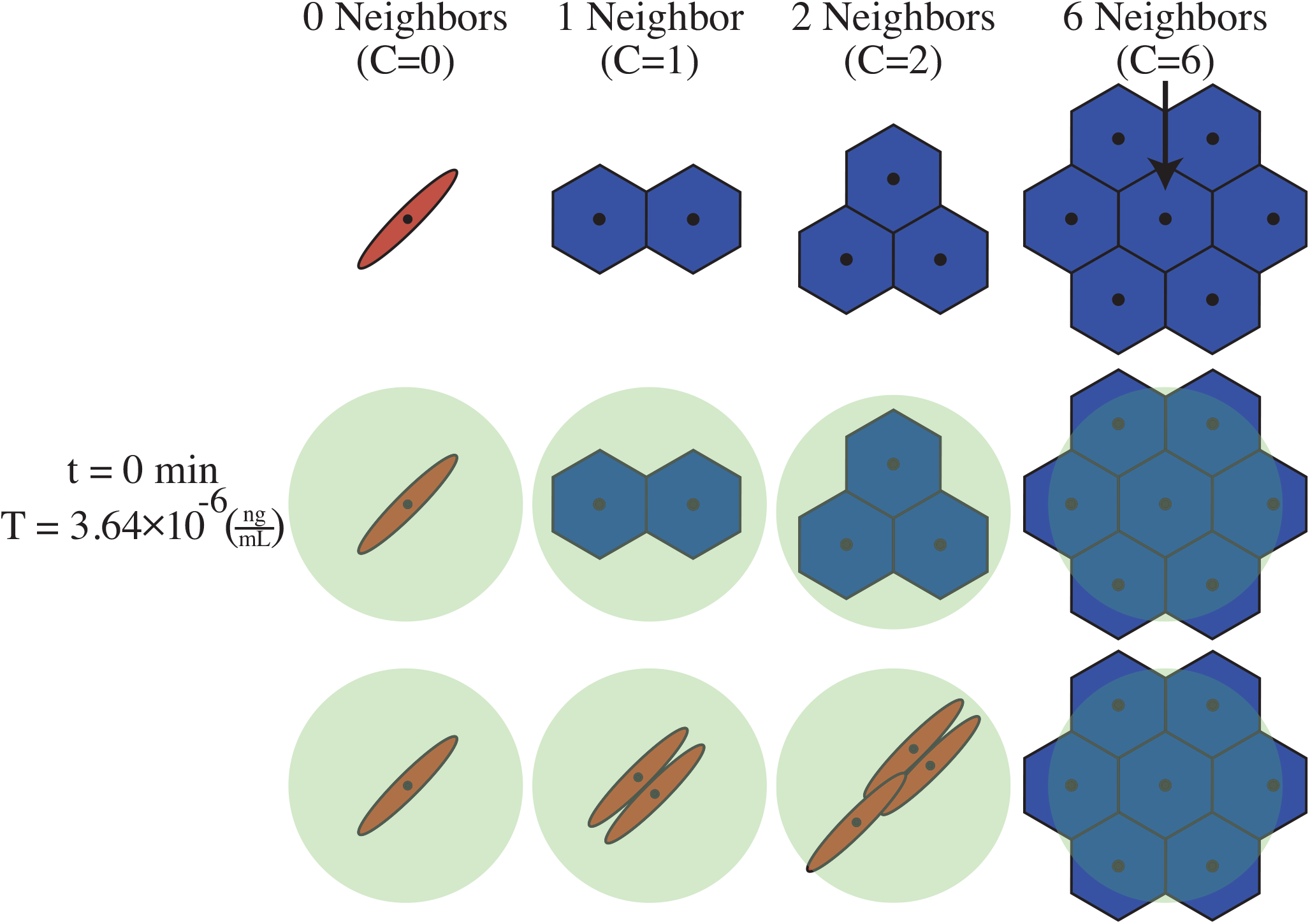
Four levels of cell-cell contact (*C* = 0, 1, 2, and 6) and two levels of exogenous TGF-*β* 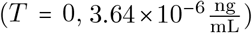 were considered for analysis. It is assumed that each cell takes up 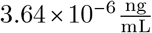 when TGF-*β* is applied. Cells without any cellular contact (*C* = 0 begins as a mesenchymal cell and exposure to exogenous TGF-*β* at *t* = 0 min maintains this phenotype. At *t* = 0 min, exogenous TGF-*β* is added to the system, allowing cells with *C* = 1, 2 neighbors to undergo EMT and adopt the mesenchymal phenotype and invasive characteristics. However, for cells with *C* = 6 neighbors, cellular contact is too high and the addition of 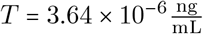 does not allow the cell to undergo the bistable switch to the mesenchymal phenotype.

### 3.2. Latin Hypercube Sampling and Partial Rank Correlation Coefficient

Latin Hypercube Sampling (LHS) and Partial Rank Correlation Coefficient (PRCC) were used to determine how sensitive the values of E-cadherin (*E*) and Slug (*S*) at *t* = 10000 min are to changes to the parameters in Equations 1-2. In the original model published by Gasior et al., steady state values were used to construct bifurcation diagrams [8]. This work uses the methodology proposed by Gasior, which allows time course simulations to run until *t* = 10000 min before recording the value of E-cadherin and Slug [9]. As shown in Supplementary Figure S1, with the parameter values put forth in Table 1, the system achieves steady state behavior prior to *t* = 50 min. However, to ensure that the long-term behavior of the system was captured under LHS-PRCC analysis, time was allowed to run to *t* = 10, 000min. For this analysis, 11 parameters {*α*_1_, *α*_2_, *β*_1_, *β*_2_, *k*_0_, *k*_1_, *k*_2_, *IC*_*S*_, *IC*_*E*_, *IC*_*T*_, *IC*_*C*_} were varied. LHS requires that the relationship 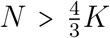 be maintained, where *K* is the number of parameters varied (*K* = 11) and *N* is the number of samples taken from each parameter range (*N* > 15) [23]. Parameter ranges were taken to be ±25% and ±50% of the value listed in Table 1. These ranges are listed in Table 2. Analysis was carried out at four different levels of cell-cell contact (*C* = 0, 1, 2, 6 cells) and two different levels of exogenous TGF-*β* 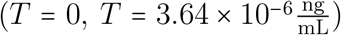. Initial conditionswere set corresponding to the level of cell-cell contact (*C*) without exogenous TGF-*β* and are detailed in Table 3.

**Table 2:** Parameter ranges for ±10%, ±25%, and ±50% of the original value in Table 1. PRCC analysis in [9] was carried out over the ±10% range. PRCC analysis in this work is carried out over the ±25% and ±50% ranges.

| Parameter | 10% Range | 25% Range | 50% Range |
| --- | --- | --- | --- |
| $\alpha_1$ | [0.1742, 0.2129] | [0.1452, 0.2419] | [0.0968, 0.2903] |
| $\alpha_2$ | [0.0772, 0.0932] | [0.0772, 0.1172] | [0.0772, 0.1572] |
| $\beta_1$ | [5.4315, 6.6385] | [4.5263, 7.5438] | [3.0175, 9.0525] |
| $\beta_2$ | [0.7470, 0.9130] | [0.6225, 1.0375] | [0.4150, 1.2450] |
| $k_0$ | [0.1120, 0.1370] | [0.0938, 0.1556] | [0.0623, 0.1868] |
| $k_1$ | [0.0669, 0.0829] | [0.0429, 0.0829] | [0.0029, 0.0829] |
| $k_2$ | [0.0360, 0.0440] | [0.0300, 0.0500] | [0.0200, 0.0600] |
| $IC_S$ | [0.0173, 0.0211] | [0.0144, 0.024] | [0.0096, 0.0288] |
| $IC_E$ | [0.0090, 0.0110] | [0.0075, 0.0125] | [0.005, 0.015] |
| $IC_T$ | $[3.2760 \times 10^{-6}, 4.0040 \times 10^{-6}]$ | $[2.73 \times 10^{-6}, 4.55 \times 10^{-6}]$ | $[1.82 \times 10^{-6}, 5.46 \times 10^{-6}]$ |
| $IC_C$ | [1.9530, 2.3870] | [1.6275, 2.7125] | [1.0850, 3.2550] |

**Table 3:**
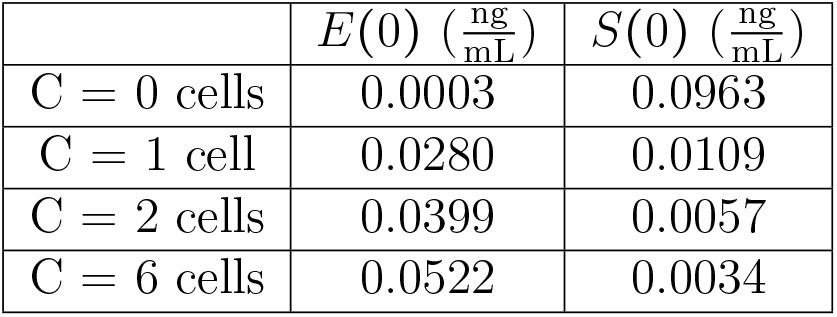
Initial conditions for different levels of cellular contact used in all PRCC analysis.

For each parameter, *N =* 10000 samples were chosen from uniform distributions across the parameter range. Time course simulations were run in MATLAB until *t* = 10000 min. The values of E-cadherin (*E*) and Slug (*S*) were recorded and rounded to the nearest 10 ^−4^ to ensure monotonicity. PRCC transformed the input parameter values and the values of *E* and *S* for each run into ranked values and measured the correlation between the rank - transformed inputs and outputs [38]. The PRCC output parameter *ρ* can range from −1 to 1 and it is standard practice to use |*ρ*| = 0.5 as a cut-off to determine significance [26]. Thus, ~*ρ*~ = 0.5 was used for sensitivity and is marked in Figures 3-4 by a dashed line. PRCC between the 11 parameters under all 8 treatment groups revealed |*ρ*_*ij*_| < 0.5∀*i* ≠ *j =* 1, …, 11, indicating that the parameters of Equations 1 and 2 are not correlated. Code available upon request.

**Figure 3:**
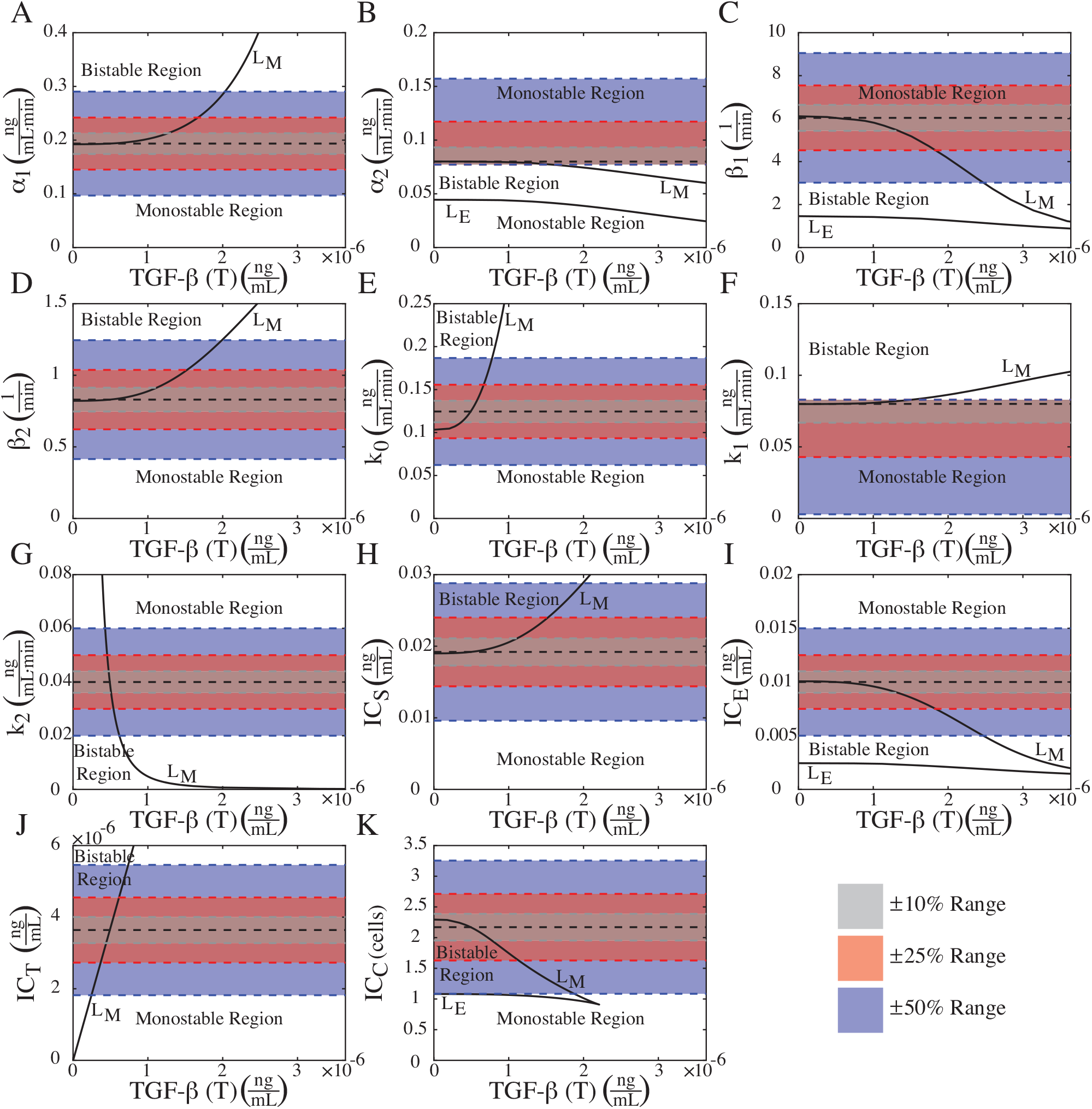
Two parameter bifurcation diagrams for TGF-*β* when a cell has 1 neighbor (*C* 1). The original parameter value from Table 1 is marked with a black dashed line. Without 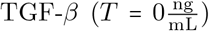, the cell begins in the bistable region in the epithelial state. If the amount of exogenous TGF-*β* exceeds the necessary threshold (*L*_*M*_ *=* 5.0224 × 10 ^−7^), the cell transitions to the mesenchymal state and enters the monostable region. If the level of exogenous TGF-*β* is reduced back to *T* = 0, the cell does not cross the *L*_*E*_ threshold and retains the mesenchymal phenotype. The parameter ranges used for PRCC analysis are marked in gray (±10%, [9]), red (±25%), and blue (±50%). For parameters {*α*_1_, *β*_1_, *β*_2_, *k*_0_, *IC*_*S*_, *IC*_*E*_, *IC*_*C*_}, the ranges cover roughly equal amounts of the bistable and monostable regions when 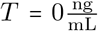. For *k* and *IC*_*T*_, the size of the bistable region only allows for the three parameter ranges to capture values in the bistable region. Conversely, the skewing of the parameter ranges for *α*_2_ and *k*_1_ show that only a small portion of values from the bistable region are sampled when 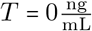. For all parameters, when 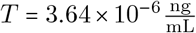, all three ranges capture only the original monostable region.

PRCC requires that all parameters have a monotonic relationship with given output values [23]. The monotonic behavior for all 11 parameters across all 8 treatment groups for both parameter ranges are shown in Figures S9-S16. Note that the parameter ranges for *α*_2_ and *k*_1_ in Table 2 have been shifted from ±25% and ±50% of the original value in Table 1. While the overall size of each parameter range is maintained, the ranges are no longer centered at the original value due to issues with monotonic behavior and negative Slug values when varied individually.

### 3.3. Bifurcation Existence in the Original Model and with a Latin Hypercube Sampling

The model created by Gasior et al. is based on earlier experimental work of the activation of EMT in MCF7 breast carcinoma cells [8; 14]. In addition to activating EMT in *in vitro* cell cultures via exposure to exogenous TGF-*β*, the authors noted that cells without TGF-*β* and no cellular contact (*T =* 0, *C =* 0) were capable of expressing the spindle-like mesenchymal phenotype. Thus, the model proposes two bistable switches underlying EMT: a reversible switch in response to cell-cell contact (*C*) and an irreversible bistable switch with respect to exogenous TGF-*β* (*T*).

In the absence of TGF-*β* (*T =* 0), the loss of cell-cell contact (*C*) by an epithelial cell would allow it to transition to the mesenchymal state [8; 14]. But, if the mesenchymal cell was able to acquire cell-cell contact, it could transition back to the epithelial phenotype, creating a reversible bistable switch.

This model also presents an irreversible switch with respect to exogenous TGF-*β*. As shown in Figure 2, a cell without any cell-cell contact (*C =* 0) begins as a mesenchymal cell and remain a mesenchymal cell with the addition of exogenous TGF-*β*. However, for a cell with *C* = 1, 2 neighbors, if exogenous TGF-*β* is added to the cell 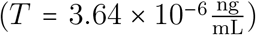, an irreversible bistable switch occurs [8]. Once the cell transitions to the mesenchymal phenotype, it is incapable of transitioning back to the epithelial phenotype, even if the exogenous TGF-*β* signal is lost, allowing it to maintain the mesenchymal phenotype. However, if the cell has 6 neighbors (*C =* 6), it begins as an epithelial cell and is incapable of undergoing the switch when exogenous TGF-*β* is added. The changes in E-cadherin and Slug with respect to TGF-*β* and the irreversible bistable behavior when *C =* 1, 2 cells, as originally shown in [9] is included as Figure S2.

The parameter values that give rise to the bistable behavior were taken from a robust bistable region. Figures 3 & 4 show the two-parameter bifurcatioshn diagrams with respect to TGF-*β* when *C =* 1 and *C =* 2 cells. The original parameter value from Table 1 is marked with a black dashed line. In the bistable region, the cell begins in the epithelial phenotype for both levels of cellular contact. Increasing TGF-*β* past the threshold of 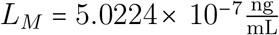 for *C* = 1 cell (Figure 3) and 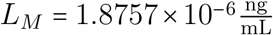 for *C* = 2 cells (Figure 4) allows the cell to transition to the mesenchymal phenotype as it enters the monostable region. If the exposure to exogenous TGF-*β* is subsequently lost 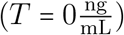, the cell will fail to cross the *L*_*E*_ threshold and will remain in the mesenchymal phenotype as it migrates away from the primary tumor. Bifurcation diagrams in Figures S2-S8 and Figures 3 & 4 were generated using XPPAUT. All code available upon request.

**Figure 4:**
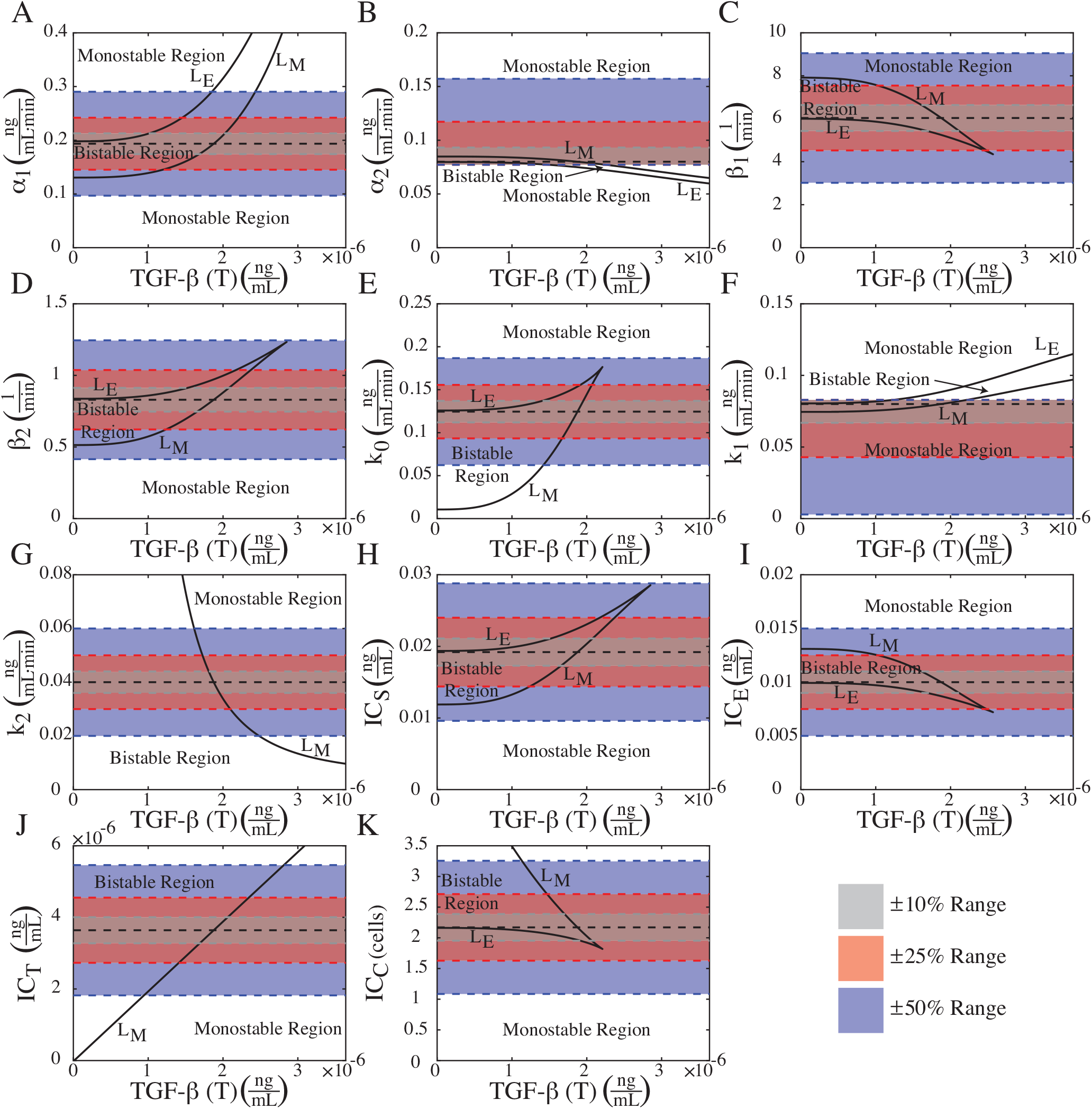
Two parameter bifurcation diagrams for TGF-*β* when a cell has 2 neighbors (*C =* 2). The original parameter value from Table 1 is marked with a black dashed line. Here, with this original value, there is still an irreverisble bistable switch with respect to TGF-*β*, but the bistable region has shifted and narrowed for parameters {*α*_1_, *α*_2_, *β*_1_, *β*_2_, *k*_0_, *k*_1_, *IC*_*S*_, *IC*_*E*_}. Now, the ±10%, ±25% and ±50% parameter ranges for PRCC analysis capture the entire bistable region when 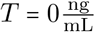. For the *IC*_*C*_ parameter, the range shifted, meaning that higher *IC*_*C*_ values used for analysis will be drawn from the original bistable region. Additionally, the ranges for *k*_2_ and *IC*_*T*_ never leave the bistable region when 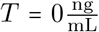. When 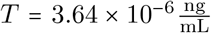, all three parameter ranges for PRCC analysis only capture the original monostable region for all parameters.

Latin Hypercube Sampling (LHS) was used to determine whether the bistable switch with respect to TGF-*β* could be preserved if all 11 parameters ({*α*_1_, *α*_2_, *β*_1_, *β*_2_, *k*_0_, *k*_1_, *k*_2_, *IC*_*S*_, *IC*_*E*_, *IC*_*T*_, *IC*_*C*_}) were varied, not just one-at-a-time. Parameter ranges were taken to be ±10%, ±25%, and ±50% of the value listed in Table 1. These ranges are listed in Table 2. Values for LHS were taken from a uniform distribution of each parameter range with *N =* 10000. Each run from the Latin Hypercube was used in Equations 1-2 and subjected to four different levels of cellular contact (*C* = 0, 1, 2, 6 cells).

Next, it is necessary to determine if LHS preserves the irreversible bistable switch with respect to TGF-*β* in this model of MCF7 cells. Previous models have examined the loss of multisability in models of EMT. In those works, the models examined the existence of an intermediate steady state expressing both epithelial and mesenchymal markers. Analysis was performed by varying parameters one-at-a-time over a ±10% range and showed that the range of values of exogenous EMT-driving factors that could produce the intermediate state was fairly robust to these parameter variations [39; 40]. Here, however, global sensitivity analysis and LHS-PRCC require that all parameters are simultaneously varied. Thus, it is unclear whether the TGF-*β* switch will be maintained.

To determine if an irreversible TGF-*β* bifurcation was preserved, each run from the Latin Hypercube was used in Equations 1-2 with the initial conditions set in Table 3 and run for *t* = 10000 min without TGF-*β* 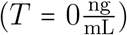. The values of E-cadherin and Slug at *t* = 10000 min (*E*_1_, *S*_1_) were then set to be the initial conditions ((*E*(*t* = 0), *S*(*t* = 0)) = (*E*_1_, *S*_2_)) for the parameter set and the value of exogenous TGF-*β* was increased to 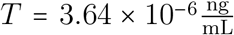 and the model was run to *t* = 10000 min. The value of E-cadherin and Slug at *t* = 10, 000 min with 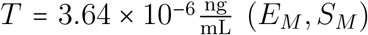 were then set as the new initial conditions ((*E*(*t* = 0), *S*(*t* = 0)) = (*E*_*M*_, *S*_*M*_) and the value of TGF-*β* was reduced to 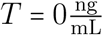 to mimic the loss of an exogenous signal. The system was run for *t* = 10000 min and the values of E-cadherin and Slug were taken as (*E*_2_, *S*_2_).

The percent change in E-cadherin and Slug values (*E*_*D*_, *S*_*D*_) before and after TGF-*β* (application and loss) was calculated using Equation 3-4 and percents were rounded to the nearest value. All runs took place in MATLAB. Code available upon request.

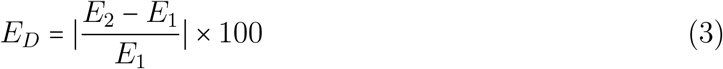

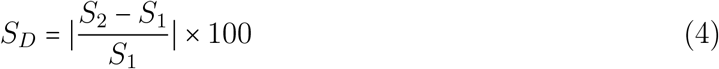

## 4. Results

### 4.1. Larger parameter ranges and tumor heterogeneity shift PRCC results

Changing the parameter range from ±25% to ±50% of the original value did not alter the PRCC results for the steady state value of E-cadherin. Instead, the most consequential factor was the level of cell-cell contact. For *C* = 0 both with and without TGF-*β*, Figures 5 and 6 show that it is the production rate of Slug (*α*_2_) and the half maximal concentration of Slug necessary for it to suppress E-cadherin transcription (*IC*_*S*_) affecting the levels of membrane-bound E-cadherin. Once the cell gains at least one neighbor (*C* ≥ 1), the parameters impacting the steady state level of E-cadherin are directly associated with E-cadherin. For cells with 1-2 neighbors (*C* = 1, 2), both with and without TGF-*β*, Figures 5C-F and 6C-F show that it is the half maximal level of cell-cell contact necessary to draw E-cadherin to the membrane for activation (*IC*_*C*_) that is influencing the amount of membrane-bound E-cadherin. Then, as the cell gains more cell-cell contact, the PRCC results shift again. For cells with *C* = 6 neighbors 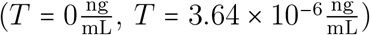, results for both sets of parameter ranges reveal that it is now the degradation rate of E-cadherin (*β*_1_) and the rate at which E-cadherin can move to the membrane to form intercellular interactions (*k*_0_) that impact the steady state value of E-cadherin.

**Figure 5:**
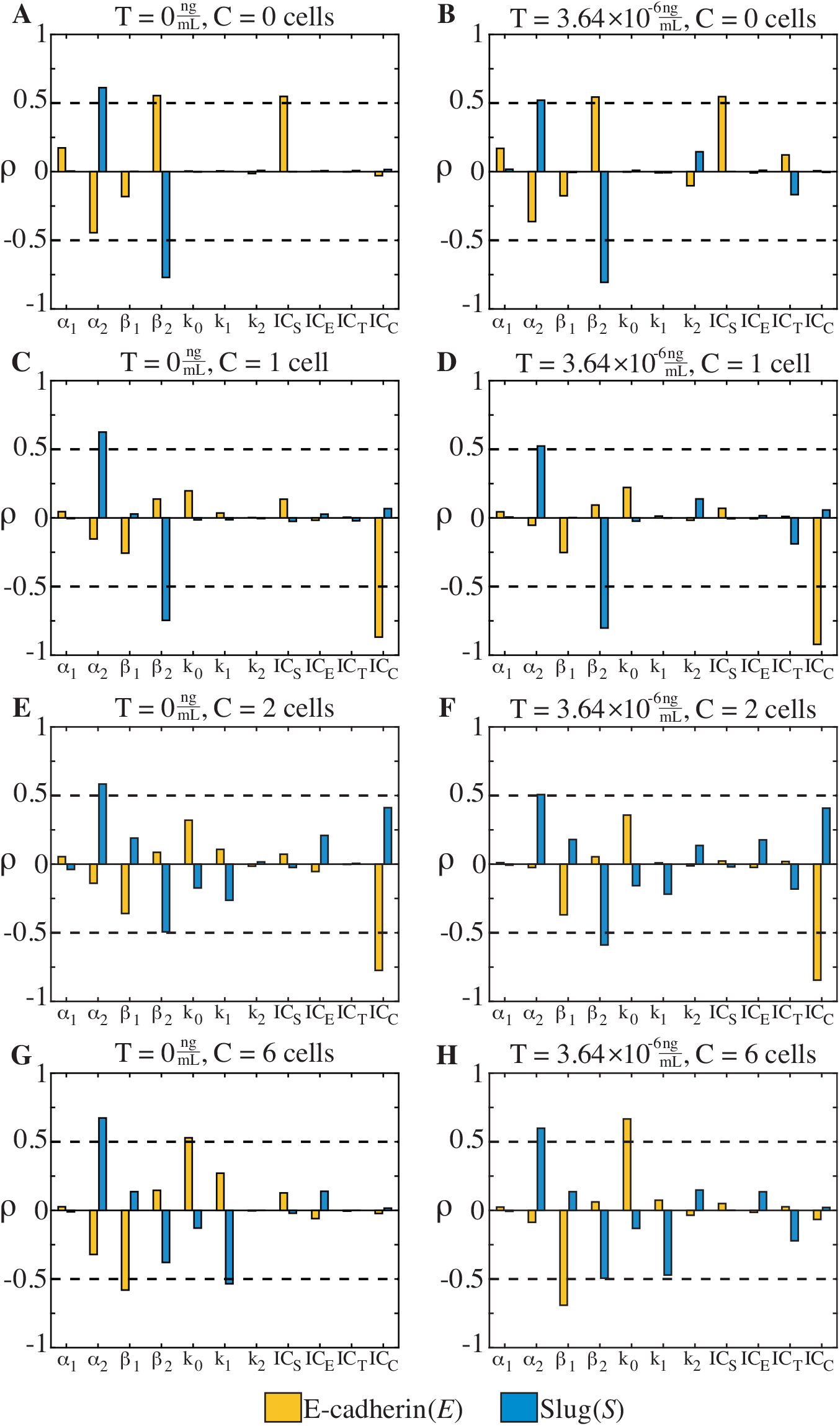
PRCC results of the 25% parameter range for the 11 parameters where the QOIs are the steady state values of E-cadherin (*E*) and Slug (*S*). Analysis was completed for 2 levels of TGF-*β* 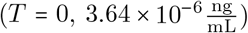 and four levels of cellular contact (*C =* 0, 1, 2, 6 cells). Cut off for significance is marked in dashed lines for each treatment group (~*ρ*~ = 0.5). If *C* = 0, E-cadherin depends on *β*_2_ and *IC*_*S*_. For *C* = 1, 2, E-cadherin is sensitive to changes in *IC*_*C*_. Increasing cell-cell contact to *C* = 6 changes the parameters impacting E-cadherin to *β*_1_ and *k*_0_. For all treatment groups, Slug (*S*) is dependent upon *α*2. Additionally, for *C* = 0, 1, 2, Slug is sensitive to changes in *β*_2_. Once *C* = 6, for 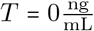, Slug depends on *k*1. For *C* = 6 cells and 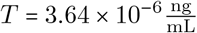, Slug is impacted by *α*_2_ and *β*_2_ using ~*ρ*~ > 0.5.

**Figure 6:**
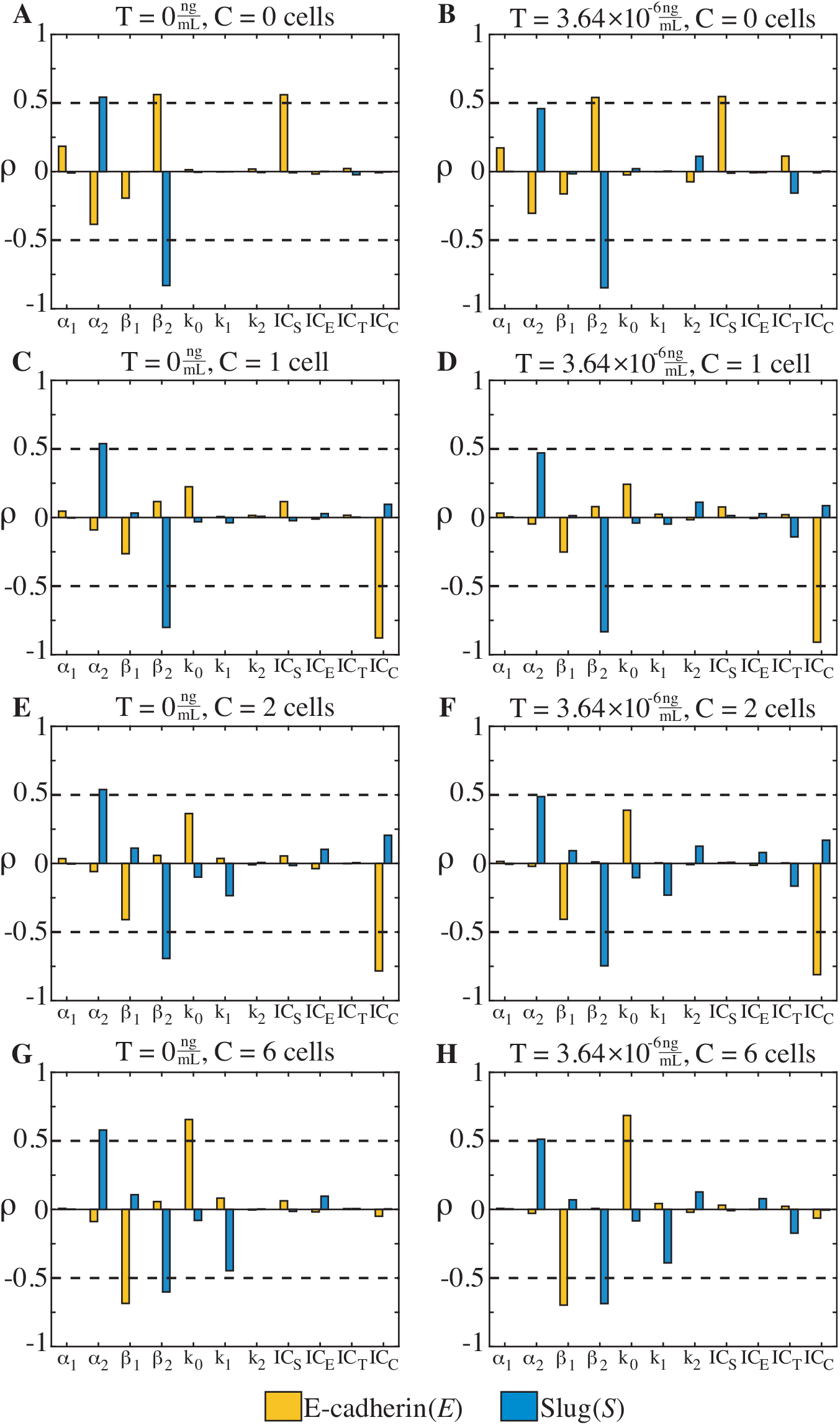
PRCC results of the ±50% parameter range for the 11 parameters where the QOIs are the steady state values of E-cadherin (*E*) and Slug (*S*). Analysis was completed for 2 levels of TGF-*β* 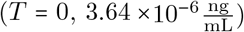 and four levels of cellular contact (*C* = 0, 1, 2, and 6 cells). Cut off for significance is marked in dashed lines for each treatment group (~*ρ*~ = 0.5). If *C* = 0, E-cadherin depends on *β*_2_ and ICS. For *C* = 1, 2, E-cadherin is sensitive to changes in *IC*_*C*_. Increasing cell-cell contact to *C* = 6 changes the parameters impacting E-cadherin to *β*_1_ and *k*_0_. For all treatment groups, Slug (*S*) depends on *β*_2_. For 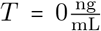 and all levels of cell-cell contact, Slug (*S*) is also dependent upon *α*_2_. For (*C, T*) = (2, 3.64 × 10^−6^) and (*C, T*) = (6, 3.64 × 10^−6^), *α*_2_ considered a significant parameter that impacts Slug using ~*ρ*~ > 0.5.

PRCC analysis for steady state value of Slug (*S*) shows that it is both the treatment groups and the parameter ranges that can impact the results. With a ±25% parameter range, Figure 5 shows the rate of Slug production (*α*_2_) is determined to significantly impact the steady state value of Slug for all levels of cell-cell contact and TGF-*β*. Additionally, for *C* = 0, 1, 2 cells, PRCC finds that the degradation rate of Slug (*β*_2_) is also important for Slug at both levels of TGF-*β*. However, for the highest level of cell-cell contact (*C* = 6 cells), without exogenous TGF-*β* 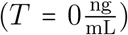, the impact of Slug degradation (*β*_2_) on the steady state value is replaced by the rate at which E-cadherin suppresses Slug activation (*k*_1_). With the addition of TGF-*β* 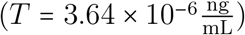 in Figure 5H, PRCC highlights the importance of Slug production and degradation (*α*_2_ and *β*_2_, respectively) using the |*ρ*| *>* 0, 5 threshold.

When the parameter range is increased to ±50% of the original value, PRCC analysis reveals new patterns of sensitivity for the steady state value of Slug. With this larger parameter range, PRCC identifies the degradation rate of Slug (*β*_2_) as the parameter that will impact the steady state of Slug across all treatment groups. Further, PRCC shows that the impact of the production rate of Slug (*α*_2_) can shift based on the level of cell-cell contact and TGF-*β*. As shown in Figure 6, when 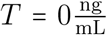, for all levels of cell-cell contact, *α*_2_ is responsible for changes in the steady state value of Slug. But, once the level of exogenous TGF-*β* is increased 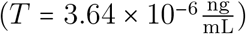, the value of *α*_2_ does not significantly impact Slug for lower levels of cell-cell contact (*C* = 0, 1 cells). But, increasing contact to *C* = 2, 6 cells, shows that the influence of *α*_2_ on Slug grows and is classified as an important parameter by PRCC.

### 4.2. Extending the parameter range for LHS reduces and eliminates the bistable switch

Figures 3 & 4 lay out the robust bifurcation region each model parameter was selected from by using two-parameter bifurcations of each parameter vs. varying input levels of TGF-*β* (*T*). The parameter ranges chosen for LHS-PRCC analysis are overlayed to show ±10% (gray), ±25% (red), and ±50% (blue) of the original value in Table 1. In Figure 3 with 1 cellular neighbor and in the absence of TGF-*β* ((*C, T*) = (1, 0)), each range covers roughly equal amounts of the bistable and monostable regions for parameters {*α*_1_, *β*_1_, *β*_2_, *k*_0_, *IC*_*S*_, *IC*_*E*_, *IC*_*C*_}. However, for other parameters, the regions are skewed. For *k*_2_ and *IC*_*T*_, the size of the bistable region only allows for the three parameter ranges to capture values in the bistable region. Additionally, the skewed parameter ranges of *α*_2_ and *k*_1_ for PRCC analysis show that only a small portion of values from the original bistable region are sampled.

If contact is increased to *C* = 2 cells, the bistable region changes. As shown in Figure 4, the bounds of the bistable region for parameters {*α*_1_, *α*_2_, *β*_1_, *β*_2_, *k*_0_, *k*_1_, *IC*_*S*_, *IC*_*E*_} have shifted, narrowing the overall area. The values of *L*_*M*_, *L*_*E*_, or both change when 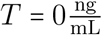. Now, the ±10%, ±25% and ±50% parameter ranges capture the entire bistable region in the absence of TGF-*β*. This is particularly different for the *α*_2_ and *k*_1_ parameter, which now have very small bistable regions that are captured by all three ranges. While the bistable region for {*k*_2_, *IC*_*T*_} grew, the ranges for *k*_2_ and *IC*_*T*_ never left the bistable region, consistent with *C* = 1. Thus, the parameter choices with 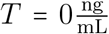 were chosen from the original bistable region for all three ranges. Finally, the bistable region for the parameter *IC*_*C*_ both shifted and expanded. Now, with (*C, T*) *=* (2, 0), higher values for analysis were drawn from the original bistable region, the opposite of what occurred with (*C, T*) = (1, 0).

When exogenous TGF-*β* is added to the system, cells with both 1-2 neighbors (*C* = 1, 2) have already transitioned to the mesenchymal state when 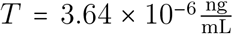. Thus, the original parameter value and all three ranges for analysis only drew from the original monostable region.

In two parameter bifurcation diagrams, such as those shown in Figures 3-4, varying only TGF-*β* while all others are fixed means that the system begins at the epithelial steady state in the bistable region with the values from Table 1 and crosses the *L*_*M*_ or *L*_*E*_ threshold that allows for the bifurcation to occur when TGF-*β* is increased. Changing the value for a given parameter in Table 1 when 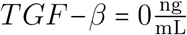 could result in beginning the system outside of the bistable region, never crossing *L*_*M*_, and thus missing this behavior altogether when TGF-*β* is increased. This is clear when two parameters are changed but harder to observe when all parameters are changed. Changing 3+ parameters could result in two possible options for the system: (1) the existence of a bistable region whose shape is altered or distorted or (2) the loss of bistable behavior altogether. A bistable region whose shape has changed could produce either an irreversible or reversible bistable switch with respect to TGF-*β*. Therefore, while the parameter ranges used by LHS-PRCC sample from both the bistable and monostable regions when 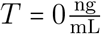 in Figures 3-4, it is unclear if LHS sampling maintains the irreversible switch with respect to TGF-*β*.

Table 4 explores the loss of an irreversible bistable switch with respect to TGF-*β* by summarizing how many of the *N* = 10000 LHS parameter sets are incapable of returning to the original steady state values at 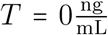 after undergoing TGF-*β* exposure for the the ±10%, ±25%, and ± 50% parameter ranges. For *C* = 1 cells under the ±10% parameter range, only 7.59% of the runs generated by the Latin Hypercube presented the possibility of an irreversible switch with respect to exogenous TGF-*β*. While both E-cadherin and Slug show > 75% difference in their values, the change in Slug is much larger. For the ±10% range, the maximum difference between E-cadherin values is 98% while the maximum difference between Slug values is 33015%. Increasing the parameter ranges ±50% meant that only 63 runs out of 10000 exhibited switch-behavior. Here, with fewer runs exhibiting the switch, the maximum E-cadherin difference is 99% and the maximum Slug difference is 8695%.

**Table 4:** Number of runs for each level of contact and parameter ranges where *E*_*D*_, *S*_*D*_ ≠ 0 and thus indicates an irreversible bistable switch. The larger the parameter range means more deviation from the true model value and fewer runs have a bifurcation. Additionally, the difference in E-cadherin values is much smaller than Slug values across all parameter ranges.

| Cells | | $\pm 10\%$ Range | | | $\pm 25\%$ Range | | | $\pm 50\%$ Range | | |
| --- | --- | --- | --- | --- | --- | --- | --- | --- | --- | --- |
|  |  | Number of Runs |  |  | Number of Runs |  |  | Number of Runs |  |  |
|  |  | > 25% | > 50% | > 75% | > 25% | > 50% | > 75% | > 25% | > 50% | > 75% |
| 0 | $E$ | 0 | 0 | 0 | 0 | 0 | 0 | 0 | 0 | 0 |
| | $S$ | 0 | 0 | 0 | 0 | 0 | 0 | 0 | 0 | 0 |
| 1 | $E$ | 759 | 759 | 759 | 195 | 195 | 195 | 63 | 63 | 62 |
| | $S$ | 759 | 759 | 759 | 195 | 195 | 195 | 63 | 63 | 63 |
| 2 | $E$ | 1280 | 1263 | 369 | 138 | 137 | 87 | 26 | 26 | 17 |
| | $S$ | 1280 | 1279 | 1277 | 138 | 138 | 138 | 26 | 26 | 26 |
| 6 | $E$ | 0 | 0 | 0 | 0 | 0 | 0 | 2 | 2 | 0 |
| | $S$ | 0 | 0 | 0 | 0 | 0 | 0 | 2 | 2 | 2 |

When contact is increased to *C* = 2, similar trends are observed as the parameter range is increased. The ±10% range has only ≈ 12.8% of the runs with the possibility of an irreversible bistable switch with > 50% difference for both E-cadherin and Slug. However, note that E-cadherin values have a smaller difference, as only 369 runs out of 10000 had > 75% difference, compared to Slug with 1277. The percent of runs with > 50% difference is decreased to 0.26% when the range widens to ±50%. But, if the difference is taken to be > 75%, the number of runs that satisfy this change for E-cadherin drops to 17.

To determine how the behavior of the switch changes when assembling the Latin Hypercube, it is necessary to look at how both E-cadherin (*E*) and Slug (*S*) change with respect to exogenous TGF-*β*, as well as changes in shape of the bistable regions from the two parameter bifurcation diagrams. Figure 7 shows three examples of runs from the Latin Hypercube sample that produced a bistable switch with respect to TGF-*β* from the ±10%, ±25%, and ±50% ranges. The parameter values used are given in Table 5. While the bifurcation is preserved, there is a key difference that is not present in either the PRCC analysis or the percent differences of Table 4: the shifting value of *L*_*M*_. For all three examples, the initial steady state values of E-cadherin and Slug are different when *C* = 1 and *C* = 2 For the ±10% range, E-cadherin and Slug begin at values of (*E, S*) = (0.0328, 0.0045) when *C* = 1 and (*E, S*) = (0.0395, 0.0022) when *C* = 2. When exogenous TGF-*β* is applied, a cell with 1 neighbor now requires at least 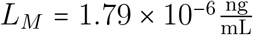 to switch to the mesenchymal phenotype while a cell with 2 neighbors requires 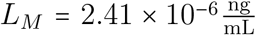. If the parameter range is increased to ±25% range, cells with *C* = 1 neighbors require 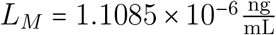 of TGF-*β* to undergo the switch. For cells with *C* = 2 neighbors, 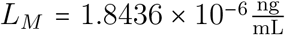 is of TGF-*β* necessary. Finally, when the parameter range is increased to ±50%, for this example, the cells now require 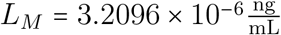 when *C* = 1 and 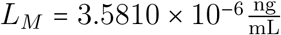 when *C* = 2. This means that the maximum value given of 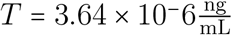 was just enough to force a transition to the mesenchymal state for *C* = 2.

**Table 5:** Parameter values for the examples shown in Figure 8. Parameter values were chosen from the ±10%, ±25%, and ±50% parameter ranges and assembled via LHS sampling. These examples produced over > 50% difference between the original steady state values of both E-cadherin and Slug with 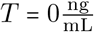 and the steady state value after TGF-*β* is applied and reduced back to 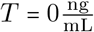.

| Parameter | $\pm 10\%$ Example<br>(8A-B) | $\pm 25\%$ Example<br>(8C-D) | $\pm 50\%$ Example<br>(8E-F) | Units |
| --- | --- | --- | --- | --- |
| $\alpha_1$ | 0.2094 | 0.1611 | 0.2325 | $\frac{\text{ng}}{\text{mL}\cdot\text{min}}$ |
| $\alpha_2$ | 0.0790 | 0.0849 | 0.0841 | $\frac{\text{ng}}{\text{mL}\cdot\text{min}}$ |
| $\beta_1$ | 6.4291 | 5.3969 | 7.4981 | $\frac{1}{\text{min}}$ |
| $\beta_2$ | 0.9010 | 1.0293 | 0.9815 | $\frac{1}{\text{min}}$ |
| $k_0$ | 0.1337 | 0.1058 | 0.0755 | $\frac{\text{ng}}{\text{mL}\cdot\text{min}}$ |
| $k_1$ | 0.0823 | 0.0819 | 0.0825 | $\frac{\text{ng}}{\text{mL}\cdot\text{min}}$ |
| $k_2$ | 0.0385 | 0.0448 | 0.0230 | $\frac{\text{ng}}{\text{mL}\cdot\text{min}}$ |
| $IC_S$ | 0.0194 | 0.0191 | 0.0260 | $\frac{\text{ng}}{\text{mL}}$ |
| $IC_E$ | 0.0103 | 0.0083 | 0.0094 | $\frac{\text{ng}}{\text{mL}}$ |
| $IC_T$ | $3.8074 \times 10^{-6}$ | $3.8572 \times 10^{-6}$ | $5.0405 \times 10^{-6}$ | $\frac{\text{ng}}{\text{mL}}$ |
| $IC_C$ | 2.3775 | 2.6806 | 3.1020 | cells |
| $n_1$ | 3 | 3 | 3 | — |
| $n_2$ | 4 | 4 | 4 | — |
| $n_3$ | 2 | 2 | 2 | — |
| $n_4$ | 3 | 3 | 3 | — |

**Figure 7:**
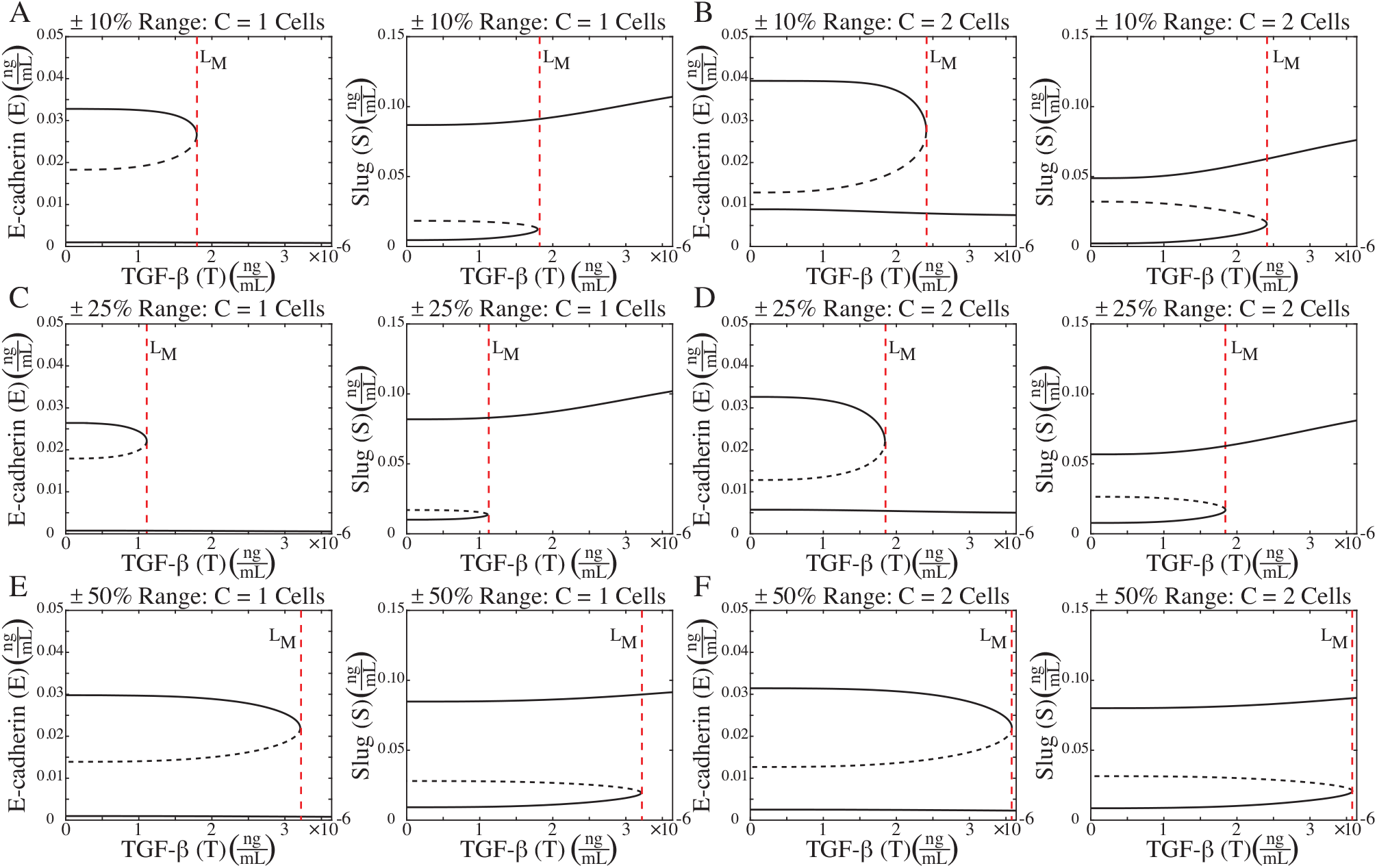
Examples of parameter sets generated from LHS using the ±10% (A&B), ±25% (C&D), and ±50% (E&F) parameter ranges. Parameter sets are listed in Table 5. These parameter sets still produce an irreversible switch with respect to TGF-*β* but the behavior has shifted. The original bistable switch occurred at 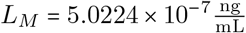 for *C* = 1 cellular neighbors and 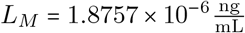 for *C* = 2 cellular neighbors [9]. The examples chosen from the LHS of the ±10% range do not switch until 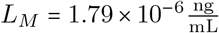 for *C* = 1 and 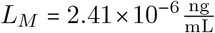 for *C* = 2. For ±25% range, this threshold changes to 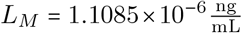 for *C* = 1 and 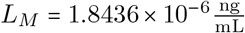 for *C* = 2 and for ±50%, this threshold increases further to 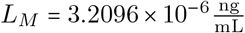 for *C* = 1 and 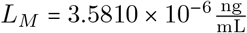 for *C* = 2.

The two parameter bifurcation diagrams for each of the cases in Figure 7 are shown in Supplementary Figures S3–S8. Like Figure 7, the overall shapes of the plots are similar to what is observed in Figures 3-4. There are both bistable and monostable regions for each parameter and the parameter values and bounds shift as the cell goes from *C =* 1 to *C =* 2 for each example. Now, however, the bistable region for each two-parameter diagram is stretched and extended. The example shown in Figure 7E-F and Supplementary Figures S7–S8 produce a large bistable region for both levels of cell-cell contact. Thus, perturbations away from the values of these systems would capture more of the bistable region when 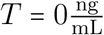. One key difference in these examples, however, is the change in the bistable region for the rate at which E-cadherin moves to the membrane due to cell-cell contact (*k*_0_). In the original model, the value presented in Table 1 was close to the *L*_*M*_ values when 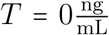. The examples in Figure 7, however, present a *k*_0_ system with a much larger bistable region. Specifically, for the ranges examined, there are not values of *k*_0_ that would force the system outside of the bistable region without TGF-*β*. This change in shape of the bistable region alters how the *k*_0_ parameter changes the phenomenological behavior of the system.

## 5. Discussion

The epithelial mesenchymal transition is a complex process spanning multiple scales in the tumor microenvironment. Exogenous pro-mesenchymal factors impact intracellular signaling cascades that control the phenotype and behavior of a given cell. Position within the tumor and exposure to factors, such as TGF-*β* can change how a cell responds to these signals. Determining how to study the impact of these exogenous factors, as well as rates associated with intracellular signaling, can present a multi-fold problem. Studying a given factor or parameter in isolation, such as with a single-parameter bifurcation diagram, can provide detailed information about how the phenomenological bistable switch changes with respect to that input. This result is informative but limited in providing detailed insight into how intracellular signaling rates affect key EMT-associated components, such as E-cadherin and Slug. Additionally, it is unclear how the impact of intracellular signaling processes changes depending on tumor placement. By turning to sensitivity analysis, such as LHS-PRCC it is possible to understand how these parameters impact the values of E-cadherin and Slug. Further, by performing this analysis under different levels of TGF-*β* and cell-cell contact acts as a method for determining some intracellular-extracellular parameter interactions. Trends emerge in the importance of certain parameters and it is possible to determine how the influence of certain intracellular processes shifts depending on tumor location.

Unfortunately, performing sensitivity analysis can be complicated by choices made early on in analysis. The process of EMT and behavioral switches underlying EMT are well studied across the literature, providing a large amount of data that can be used to help define model parameter values and ranges for analysis. This wealth of data can be incredibly beneficial when attempting to accurately capture intracellular rates and concentrations but can also be problematic if the rates vary significantly. Thus, it can be difficult to determine how large parameter ranges should be when attempting to perform analyses on models that can guide future experiments. These issues can be further complicated by the tumor microenvironment where there are multiple exogenous factors impacting intracellular signaling pathways and key EMT-associated components. The work presented here highlights the number of factors that can alter our understanding of biologically-rooted parameters when performing LHS-PRCC on a model of EMT. LHS-PRCC analysis depends on prior choices and this work shows that these choices can change the results and overall perception of a cell undergoing EMT. This is especially problematic when the transition requires a bistable switch to occur. For the given model of EMT, this work reinforces the idea that the exogenous treatment groups play a large role in determining which parameters are deemed significant, while also showing how analytical choices, such as parameter ranges, can affect the results in unknown ways.

Extending the parameter ranges to ±25% and ±50% and performing LHS-PRCC analysis produced several of the same relationships between the parameters and the steady state values of E-cadherin (*E*) and Slug (*S*) that were found in the analysis done with the ±10% range [9]. All three parameter ranges show that increasing the level of cell-cell contact and TGF-*β* shifts E-cadherin’s dependence from Slug-associated parameters (*α*_2_, *IC*_*S*_) at (*C, T*) *=* (0, 0) to E-cadherin-associated parameters (*β*_1_, *k*_0_) at (*C, T*) = (6, 3.64 × 10^−6^). Further, all three parameter ranges also highlighted the importance of *IC*_*C*_ on E-cadherin for cells with 1 or 2 neighbors (*C* = 1, 2) at both levels of TGF-*β*. For Slug, all three parameter ranges show that, for all levels of cell-cell contact when 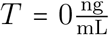, Slug depends on its own production rate (*α*_2_).

Despite these commonalities, changing the parameter range does produce different results depending upon the treatment group. While the parameters that PRCC found to be significantly impacting E-cadherin did not change as the range was increased from ±25% to ±50%, there were differences between these two ranges and the original results produced with ±10% the parameter value. For the ±10% range, E-cadherin did not have any parameters with a value of |*ρ*| *>* 0.5 for (*C, T*) *=* (0, 3.64 × 10^−6^), meaning that no conclusions could be drawn for this treatment group. Increasing the parameter range produced PRCC results that highlighted the importance of the degradation rate of Slug (*β*_2_) and the half maximal amount of Slug necessary to inhibit E-cadherin transcription (*IC*_*S*_), the same two parameters indicated for (*C, T*) *=* (0, 0). Additionally, while many of the results remain consistent, the larger parameter ranges focus on the importance of E-cadherin related parameters (*β*_1_, *k*_0_) for (*C, T*) *=* (6, 0), the original ±10% parameter range showed that it was still Slug-associated parameters (*α*_2_, *k*_1_) controlling E-cadherin for this treatment group.

Differences due to parameter ranges also occurred during LHS-PRCC analysis of Slug. Under the ±10% range, increasing cell-cell contact told a very straightforward narrative about the shift in parameters [9]. With low cell-cell contact, Slug was influenced by its own production and degradation rates (*α*_2_ and *β*_2_, respectively). Increasing cell-cell contact revealed that the importance of *β*_2_ was replaced by an increasing importance of the rate at which E-cadherin suppresses Slug (*k*_1_). Thus, increasing cell-cell contact activated E-cadherin and led to its increasing influence on Slug. However, increasing the parameter ranges shows a less clear narrative. For ±25% with *C =* 0, 1, 2 and both levels of TGF-*β*, the results of PRCC show that it is still the rates of production and degradation that control Slug (*α*_2_, *β*_2_, respectively). The rise in E-cadherin suppression (*k*_1_) does not occur until *C =* 6 and, even then, it is not as clear as it was with the smaller parameter range. Instead, for (*C, T*) = (6, 0), PRCC finds that it is now *α*_2_ and *k*_1_ impacting the steady state value of Slug but, when TGF-*β* is applied, the influence of *k*_1_ falls below the threshold for significance. Although, the *ρ* value for *β*_2_ and *k*_1_ for this treatment group are very similar, the significance threshold means that *k*_1_ is not considered important to the value of Slug. When the parameter range is increased to ±50%, PRCC no longer indicates that *k*_1_ is influencing Slug for any treatment groups. Also, the influence of *α*_2_ on Slug is also no longer guaranteed. For systems with exogenous TGF-*β* 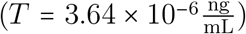, PRCC finds that *α*_2_ is not significantly impacting Slug for *C* = 0, 1. Only once there is enough cell-cell contact does *α*_2_ surpass that threshold.

Taken together, these differences in sensitivity across the three sets of parameter ranges show the potential issues that can occur when performing analysis without additional information about the system. While many of the same parameters are identified by PRCC, if the results of a specific range or treatment group was generalized to all systems, the results could be misinterpreted and could have consequences when designing future experiments focused on therapeutic intervention. For example, in the (*C, T*) = (6, 0) treatment group, the ±10% results from [9] and the ±25% results presented here suggest that the two parameters controlling Slug levels are the production rate of Slug (*α*_2_) and the rate at which E-cadherin suppresses Slug (*k*_1_). Increasing the parameter range to ±50% suggests that Slug is controlled by its production and degradation rates (*α*_2_, *β*_2_). This means a wider range of accepted parameter values change the parameters suggested as significant by PRCC, injecting doubt in a forward path.

This analysis also makes it clear that specific choices in LHS-PRCC may also play a role. This work used the threshold of |*ρ*| *>* 0.5 as a cut-off for significance and, while that cut-off has been used in the literature, it could change the understanding of important parameters for specific treatment groups from PRCC analysis [29]. For example, both the ±25% and ±50% parameter ranges, increasing the level of cell-cell contact with 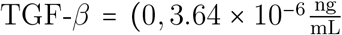 increases the influence of *β*_1_ and *k*_0_ on E-cadherin. A strict cut-off for sensitivity of ~*ρ*~ > 0.5 means that it is only when *C =* 6 cells that both parameters appear to impact E-cadherin under both parameter ranges and levels of TGF-*β*. Without this limitation, analysis would possibly have highlighted that these two parameters impact E-cadherin when *C =* 2 cellular neighbors for both parameter ranges. Similarly, the impact of the *k*_1_ parameter on Slug is missed due to the imposed |*ρ*| *>* 0.5. As shown in Figures 5-6, increasing the cell-cell contact increases the ~*ρ*~ value for *k*_1_ in both parameter ranges. However, for (*C, T*) = (6, 0), 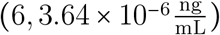 in the ±25% range and 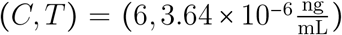 in the ±50% range, *k*_1_ never crosses the ~*ρ*~ > 0.5 threshold, meaning that its importance is missed.

While setting the value of *ρ* can affect the PRCC analysis, issues with LHS-PRCC results can also arise due to the sampling method and the construction of the Latin Hypercube. PRCC requires a monotonic relationship between the inputs and the outputs. To preserve this monotonic relationship, parameter ranges can be shifted or truncated, such as occurred with *α*_2_ and *k*_1_. Given that PRCC revealed the importance of these parameters on the steady state values of E-cadherin and Slug in different treatment groups, it is possible that this change in parameter space sampled could be impacting the results across all three ranges. Additionally, this work shows that employing LHS means that phenomenological behavior, such as the bistable switch can be lost. At best, with a small ±10% range on all parameters, ≈7.6% of runs had a large difference in E-cadherin and Slug values before and after the application and loss of exogenous TGF-*β*. As shown in the two-parameter bifurcation diagram (Figures 3 & 4), one-at-a-time sampling of each parameter could preserve the bistable switch. But simultaneously sampling all parameter ranges means that we can lose the bistability and thus lose the ability to understand how to prevent it or at what value it happens at. This is exacerbated by the unknown parameter range—large parameter ranges sample outside the original bistable region more, which then translates into fewer ranges having that switch.

This work highlights the how the chosen parameter range can affect LHS-PRCC analysis and the biological conclusions taken for future experimental development. While parameters in the literature can assist initially guiding LHS-PRCC, data variations due to caveats in its collection can inject too much doubt into the results to be beneficial in a computational-experimental feedback loop. Instead, if parameter ranges are being estimated based on literature values, performing multiple iterations of analysis with multiple range sizes is the best course of action, especially for highly complicated models with multiple inputs. Within these multiple levels of sampling, patterns that would be missed on a single iteration become clear. But it must also be considered whether LHS-PRCC are the appropriate form of analysis for phenomenological models such as this. If the researcher is only interested measurable QOIs, like the steady state values of intracellular components, LHS-PRCC can be an excellent option. Even with the caveat of monotonicity, LHS-PRCC can provide an excellent opportunity to understand which parameters affect these QOIs and whether that relationship is positive or negative. Thus, if the goal is to prevent Slug from accumulating, for example, then this methodology can pinpoint which parameters correlate with a growing Slug concentration. But, if the goal is to study phenomenological behavior as the QOI, this method may fail and employing a different form of GSA, such as Morris Method with its one-at-a-time sampling, may be more appropriate. Thus, this analysis serves to how that it is multiple conditions, including choices in the form of analysis, that produce the results shown, meaning that what is observed and learned from analysis like PRCC is as much about the conditions underpinning the model, as well as the desired QOI, as it is about what the analyst understands about the system.

## 6. Author Contributions

K. I. Gasior conceived of and carried out all the work presented here. K.I. Gasior also wrote the entirety of this manuscript.

## 7. Acknowledgements

The author would like to thank Dr. Nick Cogan for offering feedback on this manuscript.

## 8. Competing Interests

The author declares that she has no competing interests.

## 9. Data Availability

Data sharing is not applicable to this article as no datasets were generated or analysed during the current study.

## 10. Code Availability

All code used in this work will be made available upon reasonable request.

## 11. Funding

No funds, grants, or other support was received for this study or preparation of this manuscript. The author has no relevant financial or non-financial interests to disclose.

## 12. Supplementary Material

PRCC data from [9] for the ±10% parameter ranges is shown in Table S1.

Figure S2 shows the original bifurcation diagrams of the model in Equations 1-2 with respect to TGF-*β* as shown in [9]. For *C =* 0, 6, no bifurcation occurs. For *C =* 0, the cell begins as a mesenchymal cell and with the application of TGF-*β*, it remains a mesenchymal cell. For *C* = 6 cells, the cell has too many neighbors. With the application of TGF-*β*, the level of E-cadherin lowers and the amount of Slug increases but not enough to trigger a switch. For *C* = 1, 2, the cell can undergo the bistable switch with respect to TGF-*β*.

PRCC data from [9] for the ±10% parameter ranges is shown in Table S1.

Monotonicity plots for the steady state value of E-cadherin (*E*) and Slug (*S*) with respect to parameters from Equations 1-2 for both the ±25% parameter range (S9-S12) and the ±50% parameter range (S13-S16). All parameter ranges are detailed in Table 2. For each figure, the cell–cell contact is varied by column (*C =* 0, 1, 2, 6 cells) while each individual subfigure has 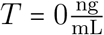 in black and 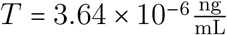 in red.

**Figure S1:**
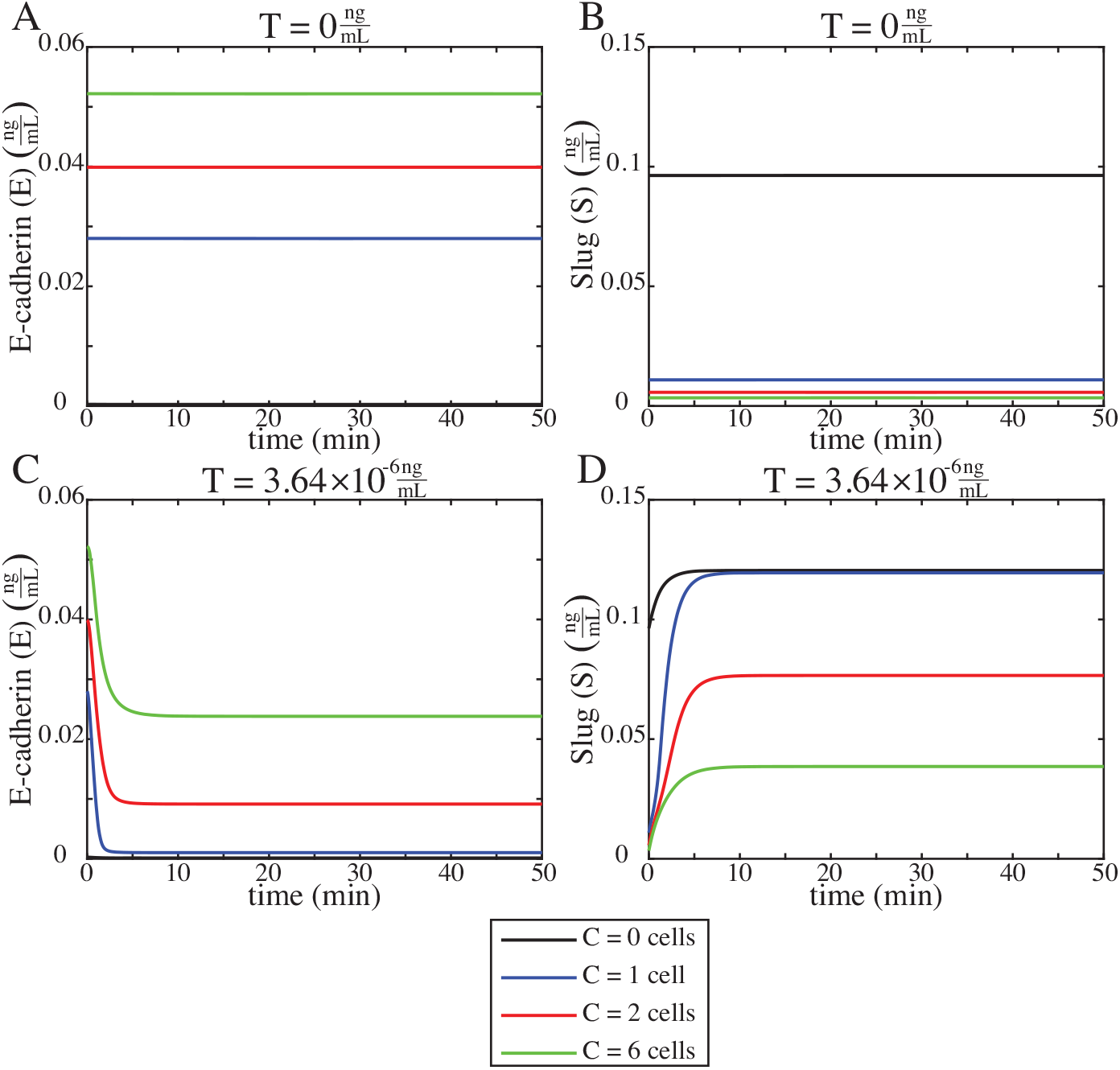
Time course simulations of E-cadherin (E) and Slug (S) from the original model in Equations 1-2 and parameter values from Table 1 when the cell is exposed to 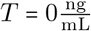 of TGF-*β* and 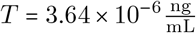 of TGF-*β*. Under these conditions, the system reaches its steady state value prior to *t* = 50min but, to ensure that the long-term behavior of the system was captured under LHS-PRCC analysis, time was allowed to run to *t =* 10, 000min.

**Figure S2:**
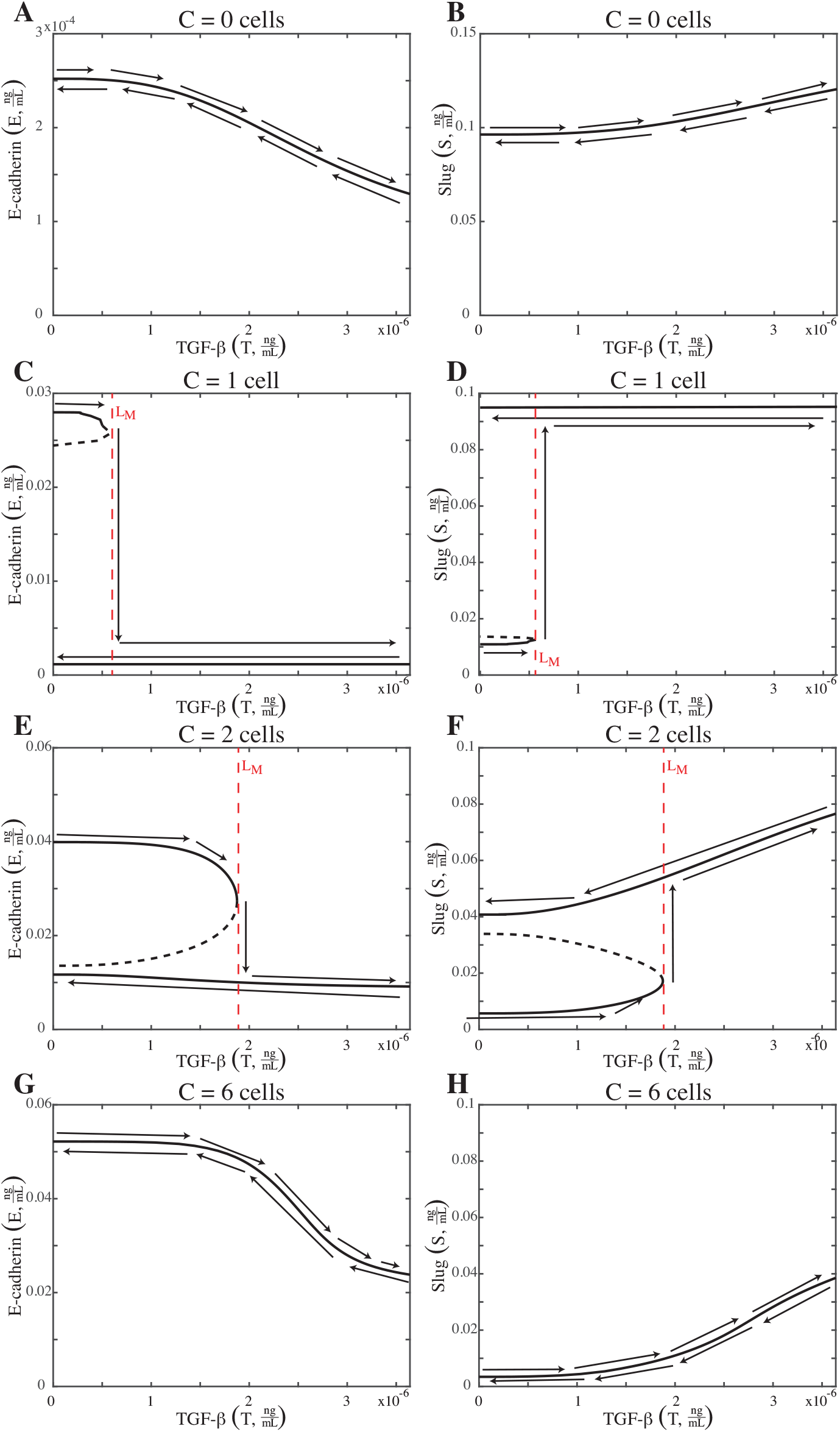
The original bifurcation diagrams of the model in Equations 1-2 with respect to TGF-*β* as shown in [9]. For *C* = 0, 6, no bifurcation occurs. For *C* = 0, the cell begins as a mesenchymal cell and with the application of TGF-*β*, it remains a mesenchymal cell. For *C* = 6 cells, the cell has too many neighbors. With the application of TGF-*β*, the level of E-cadherin lowers and the amount of Slug increases but not enough to trigger a switch. For *C* = 1, 2, the cell can undergo the bistable switch with respect to TGF-*β*. For *C* = 1 neighbors, a cell requires 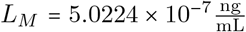 TGF-*β* to transition. For *C* = 2 neighbors, a cell requires 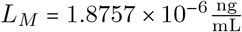 TGF-*β* to transition.

**Figure S3:**
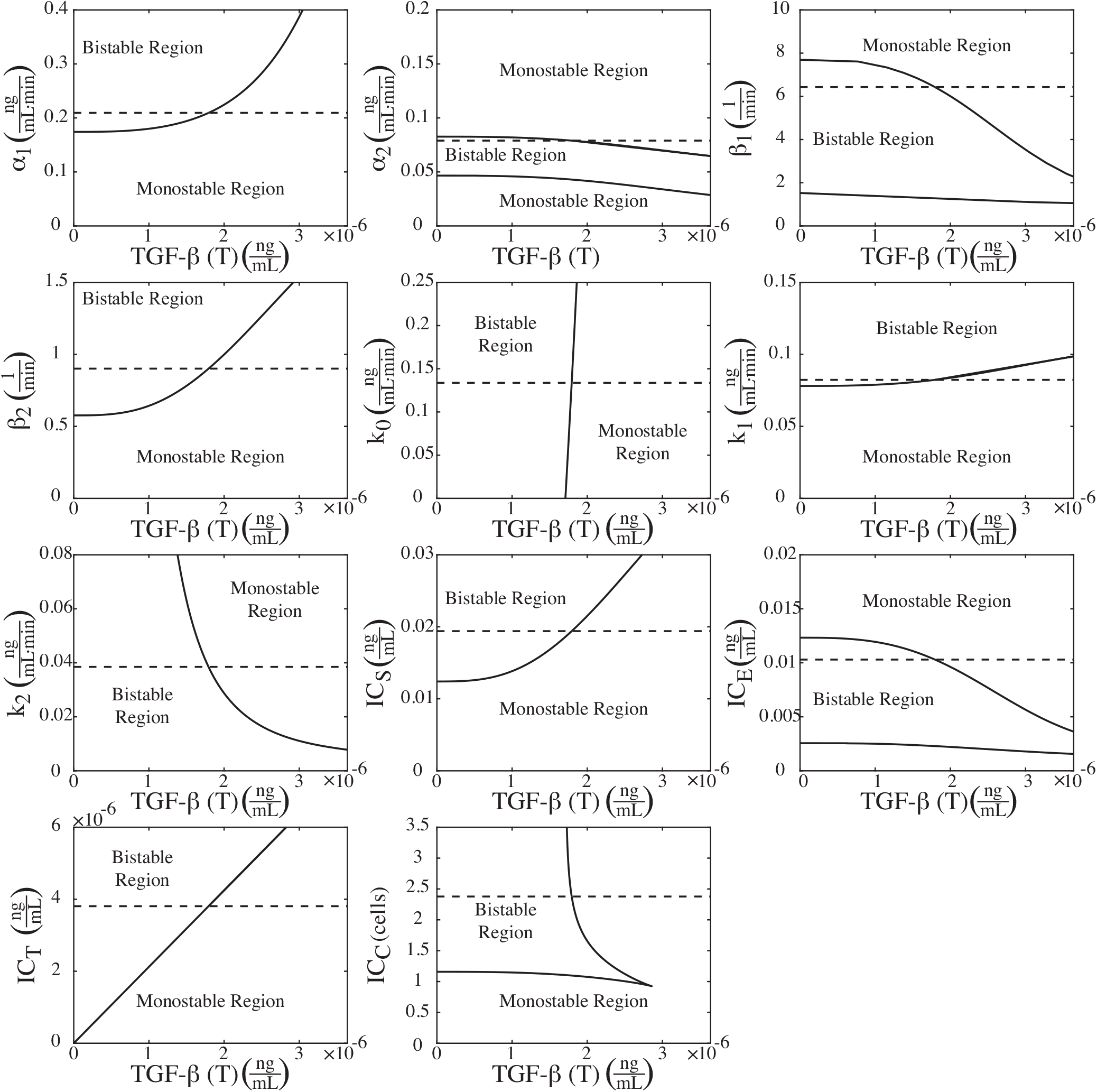
Two parameter bifurcation diagrams for the example parameter set from the ±10% range shown in Figure 7A. Here, the cell has 1 neighbor (*C* = 1). The parameter values for this example are listed in Table 5 and are shown with a dashed line.

**Figure S4:**
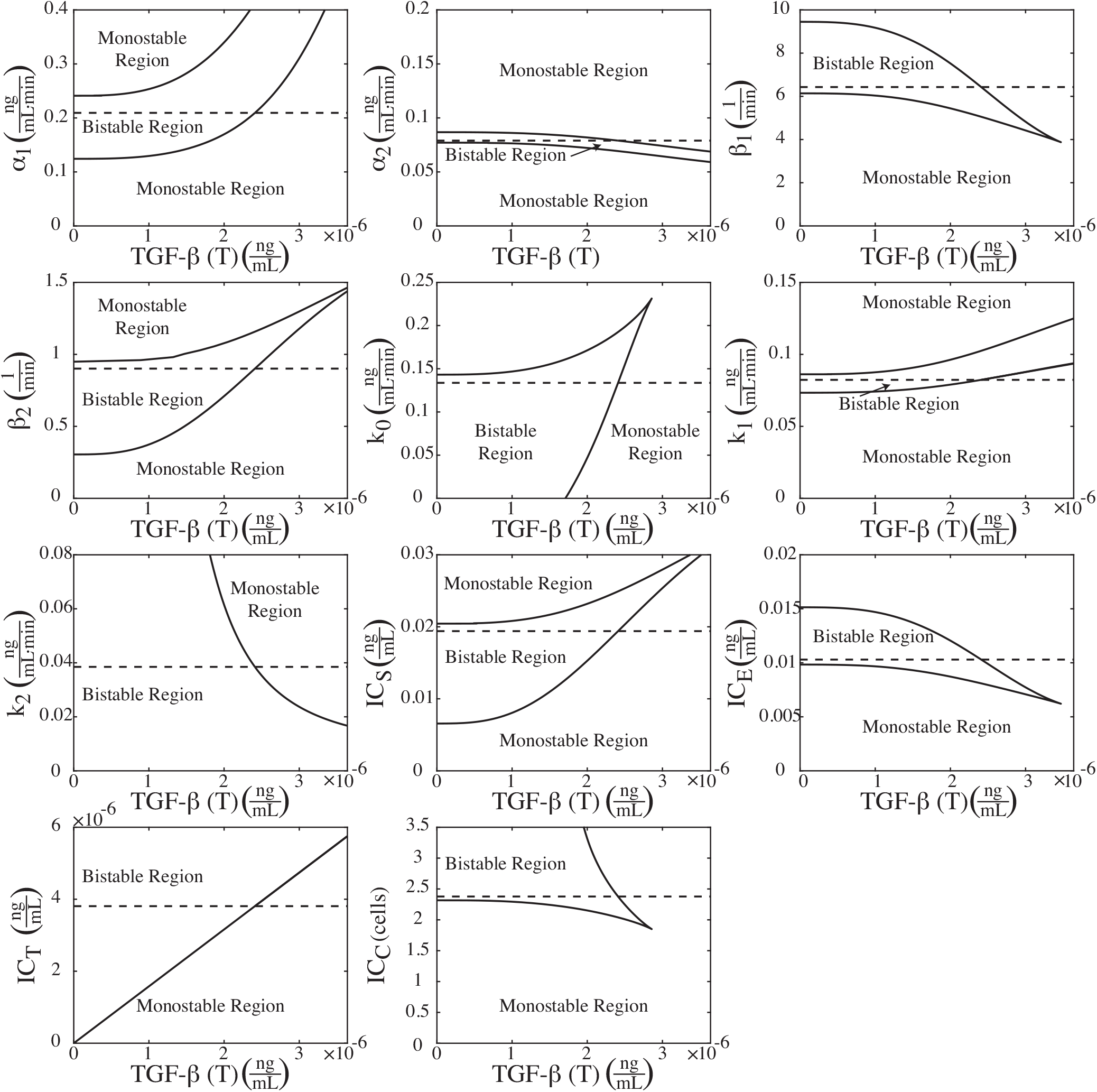
Two parameter bifurcation diagrams for the example parameter set from the ±10% range shown in Figure 7B. Here, the cell has 2 neighbors (*C* = 2). The parameter values for this example are listed in Table 5 and are shown with a dashed line.

**Figure S5:**
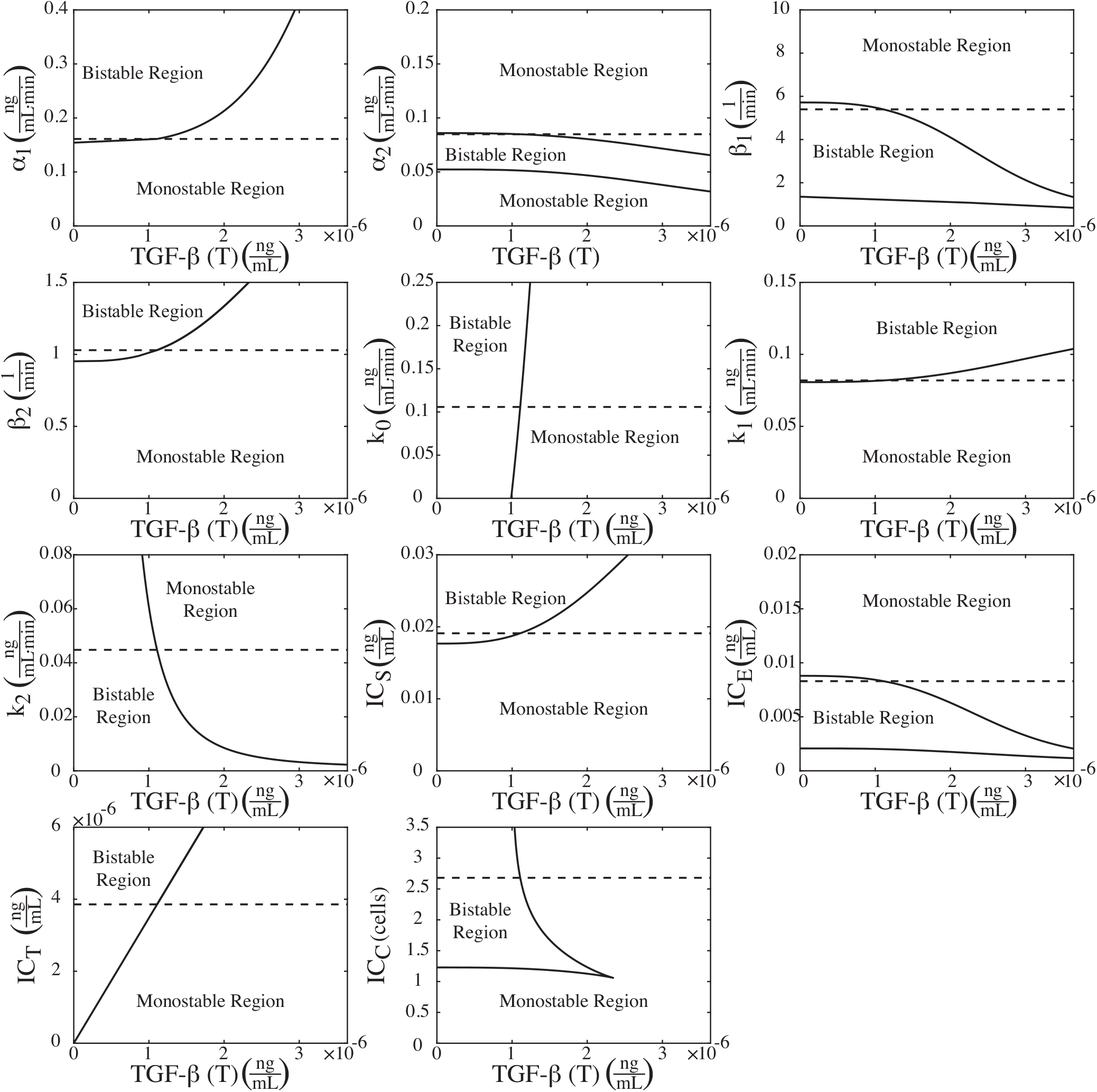
Two parameter bifurcation diagrams for the example parameter set from the ±25% range shown in Figure 7C. Here, the cell has 1 neighbor (*C* = 1). The parameter values for this example are listed in Table 5 and are shown with a dashed line.

**Figure S6:**
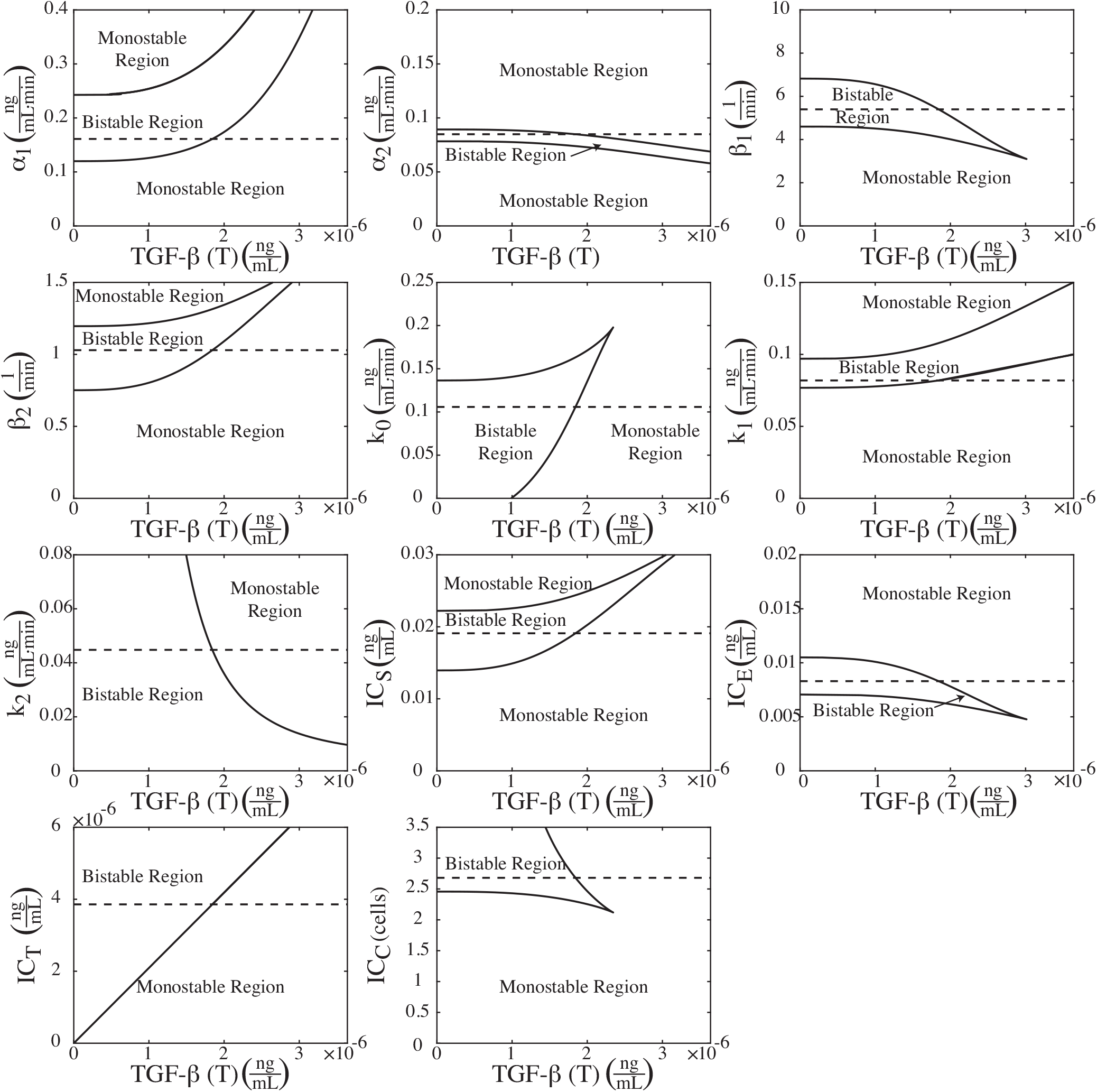
Two parameter bifurcation diagrams for the example parameter set from the ±25% range shown in Figure 7D. Here, the cell has 2 neighbors (*C* = 2). The parameter values for this example are listed in Table 5 and are shown with a dashed line.

**Figure S7:**
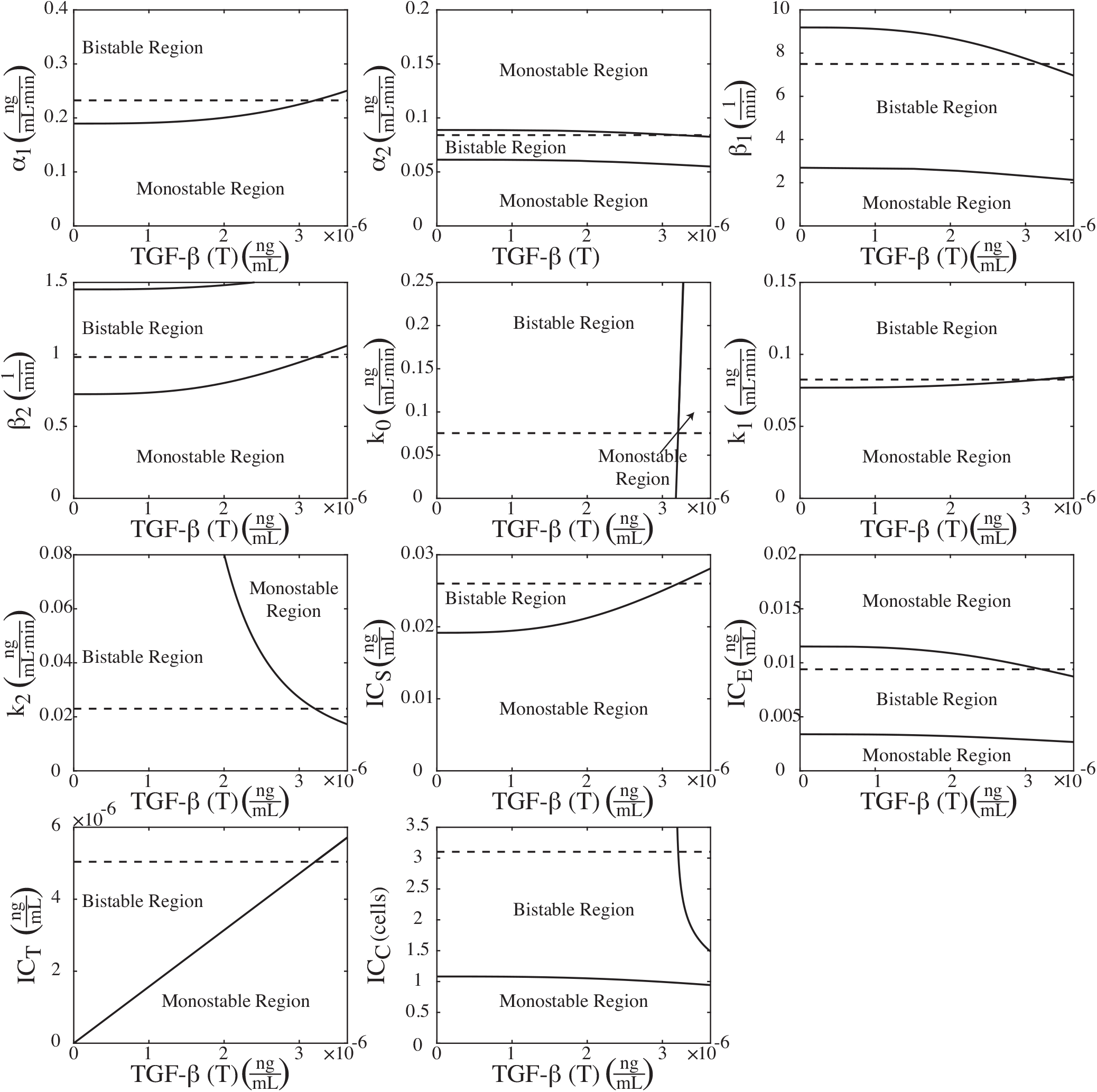
Two parameter bifurcation diagrams for the example parameter set from the ±50% range shown in Figure 7E. Here, the cell has 1 neighbor (*C* = 1). The parameter values for this example are listed in Table 5 and are shown with a dashed line.

**Figure S8:**
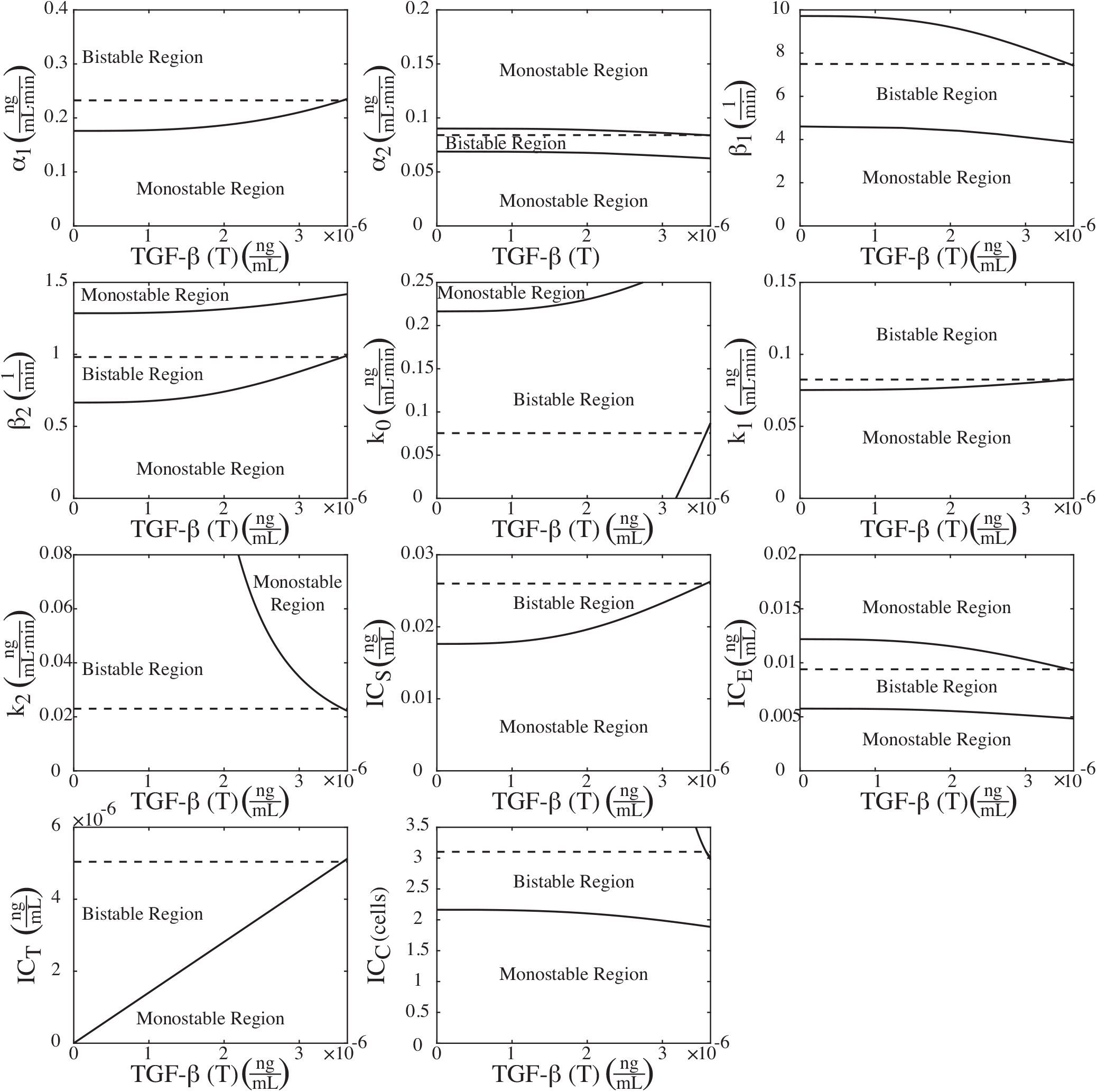
Two parameter bifurcation diagrams for the example parameter set from the ±50% range shown in Figure 7F. Here, the cell has 2 neighbors (*C* = 2). The parameter values for this example are listed in Table 5 and are shown with a dashed line.

**Figure S9:**
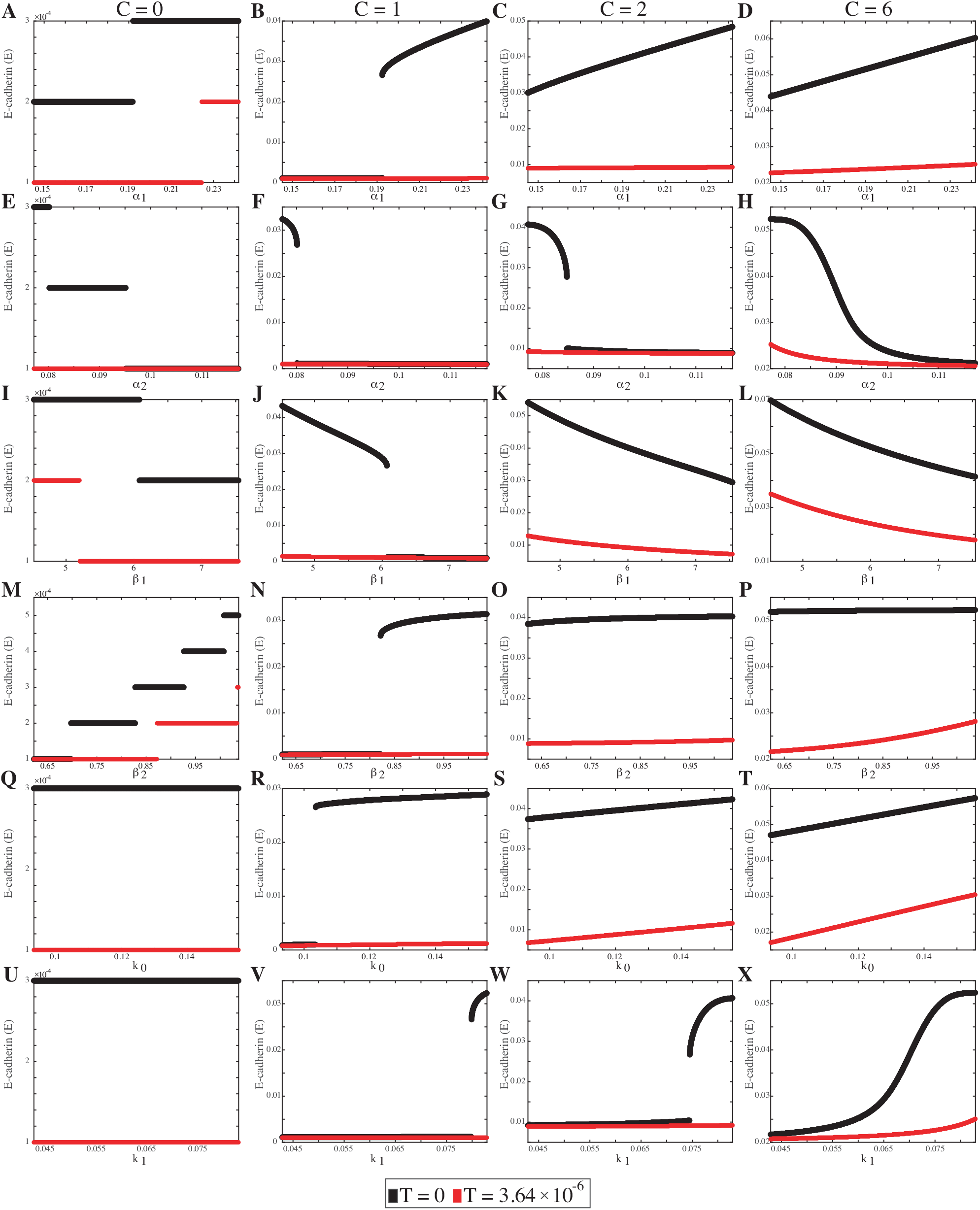
E-cadherin (E) monotonicity plots for parameters (A–D) *α*_1_, (E–H) *α*_2_, (I–L) *β*_1_, (M–P) *β*_2_, (Q–T) *k*_0_, and (U–X) *k*_1_ from Equations 1-2 with a ±25% parameter range. For each plot, 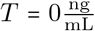 is in black and and 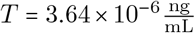 is in red. The level of cell–cell contact is given for each column with (A, E, I, M, Q, U) *C* = 0 cells, (B, F, J, N, R, V) *C* = 1 cells, (C, G, K, O, S, W) *C* = 2 cells, and (D, H, L, P, T, X) C = 6 cells.

**Figure S10:**
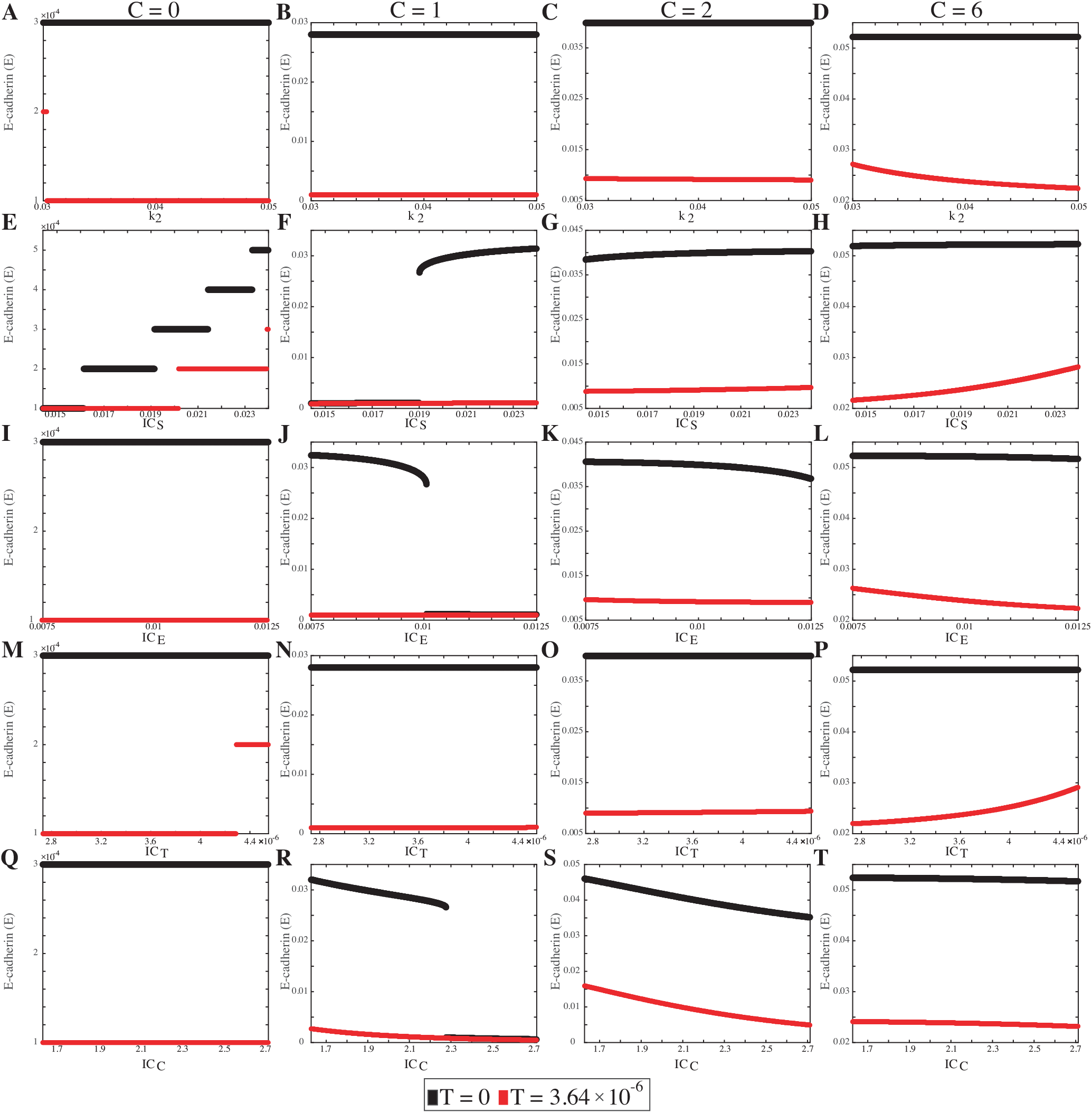
E-cadherin (E) monotonicity plots for parameters (A–D) k2, (E–H) ICS, (I–L) ICE, (M–P) ICT, and (Q–T) ICC from Equations 1-2 with a ±25% parameter range. For each plot, 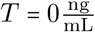 is in black and and 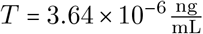 is in red. The level of cell–cell contact is given for each column with (A, E, I, M, Q) *C* = 0 cells, (B, F, J, N, R) *C* = 1 cells, (C, G, K, O, S) *C* = 2 cells, and (D, H, L, P, T) *C* = 6 cells.

**Figure S11:**
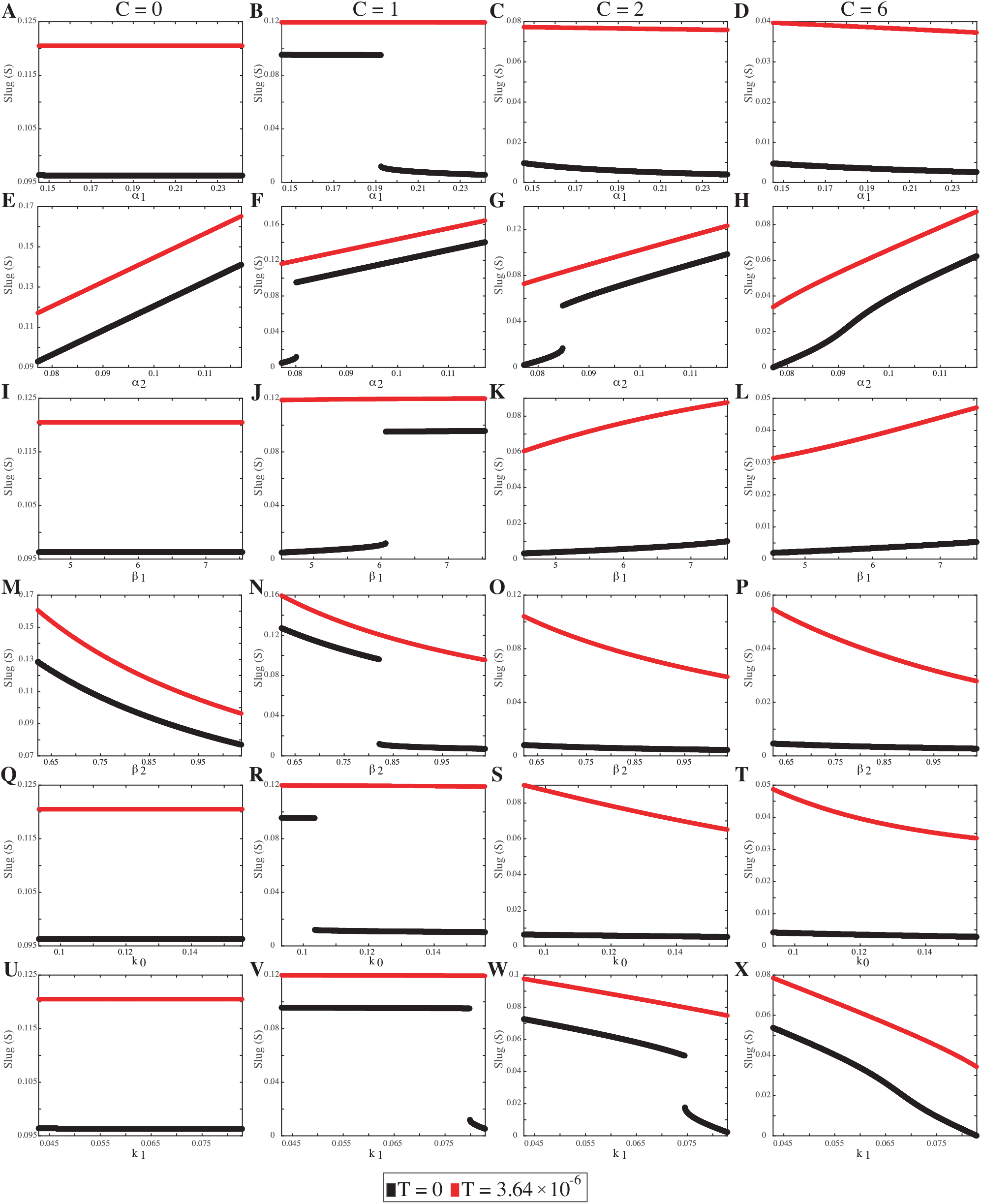
Slug (*S*) monotonicity plots for parameters (A–D) *α*_1_, (E–H) *α*_2_, (I–L) *β*_1_, (M–P) *β*_2_, (Q–T) *k*_0_, and (U–X) *k*_1_ from Equations 1-2 with a ±25% parameter range. For each plot, 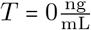 is in black and and 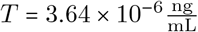 is in red. The level of cell–cell contact is given for each column with (A, E, I, M, Q, U) *C* = 0 cells, (B, F, J, N, R, V) *C* = 1 cells, (C, G, K, O, S, W) *C* = 2 cells, and (D, H, L, P, T, X) *C* = 6 cells.

**Figure S12:**
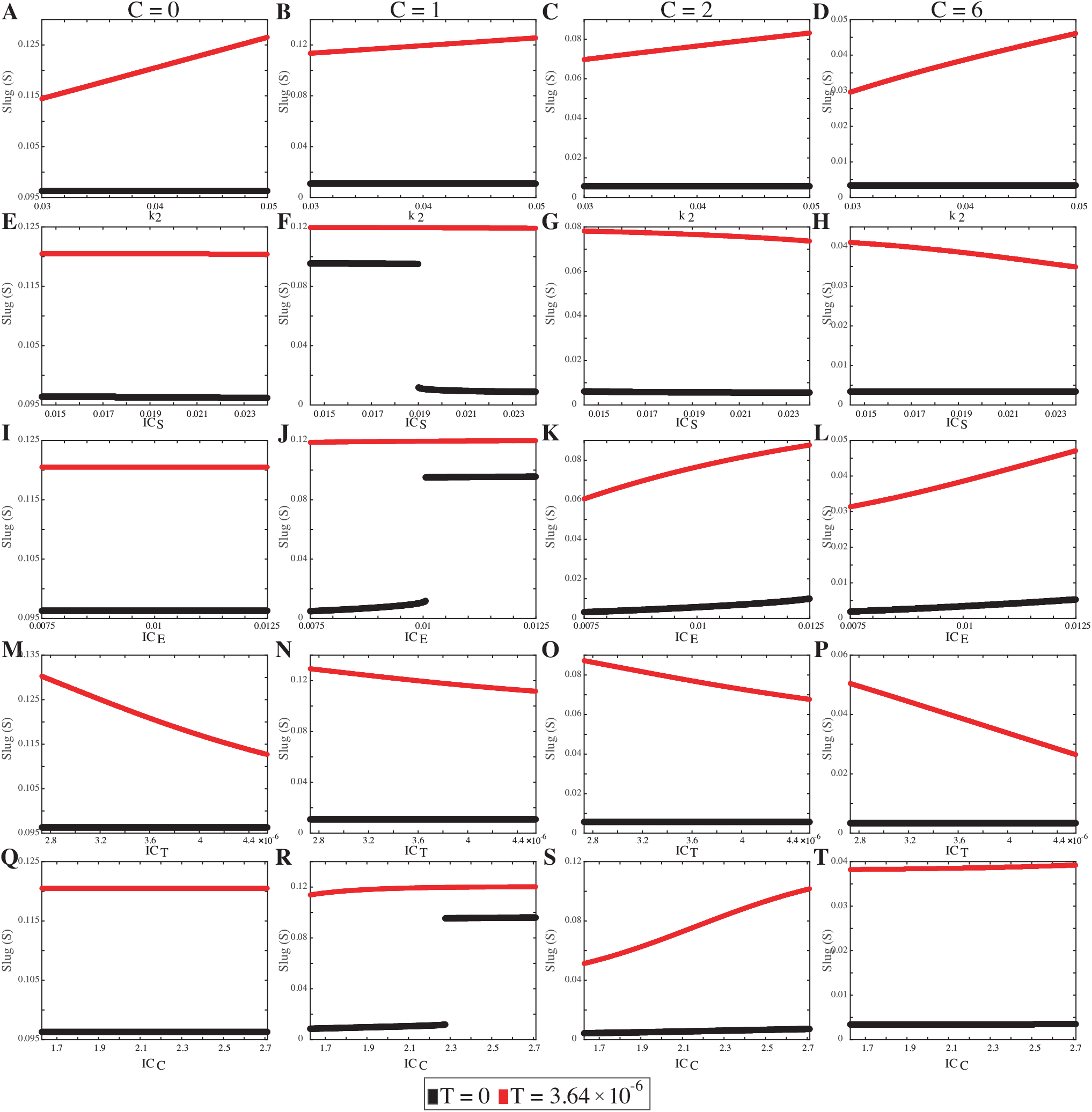
Slug (S) monotonicity plots for parameters (A–D) *k*_2_, (E–H) *IC*_*S*_, (I–L) *IC*_*E*_, (M–P) *IC*_*T*_, and (Q–T) *IC*_*C*_ from Equations 1-2 with a ±25% parameter range. For each plot, 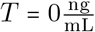 is in black and and 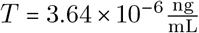 is in red. The level of cell–cell contact is given for each column with (A, E, I, M, Q) *C* = 0 cells, (B, F, J, N, R) *C* = 1 cells, (C, G, K, O, S) *C* = 2 cells, and (D, H, L, P, T) *C* = 6 cells.

**Figure S13:**
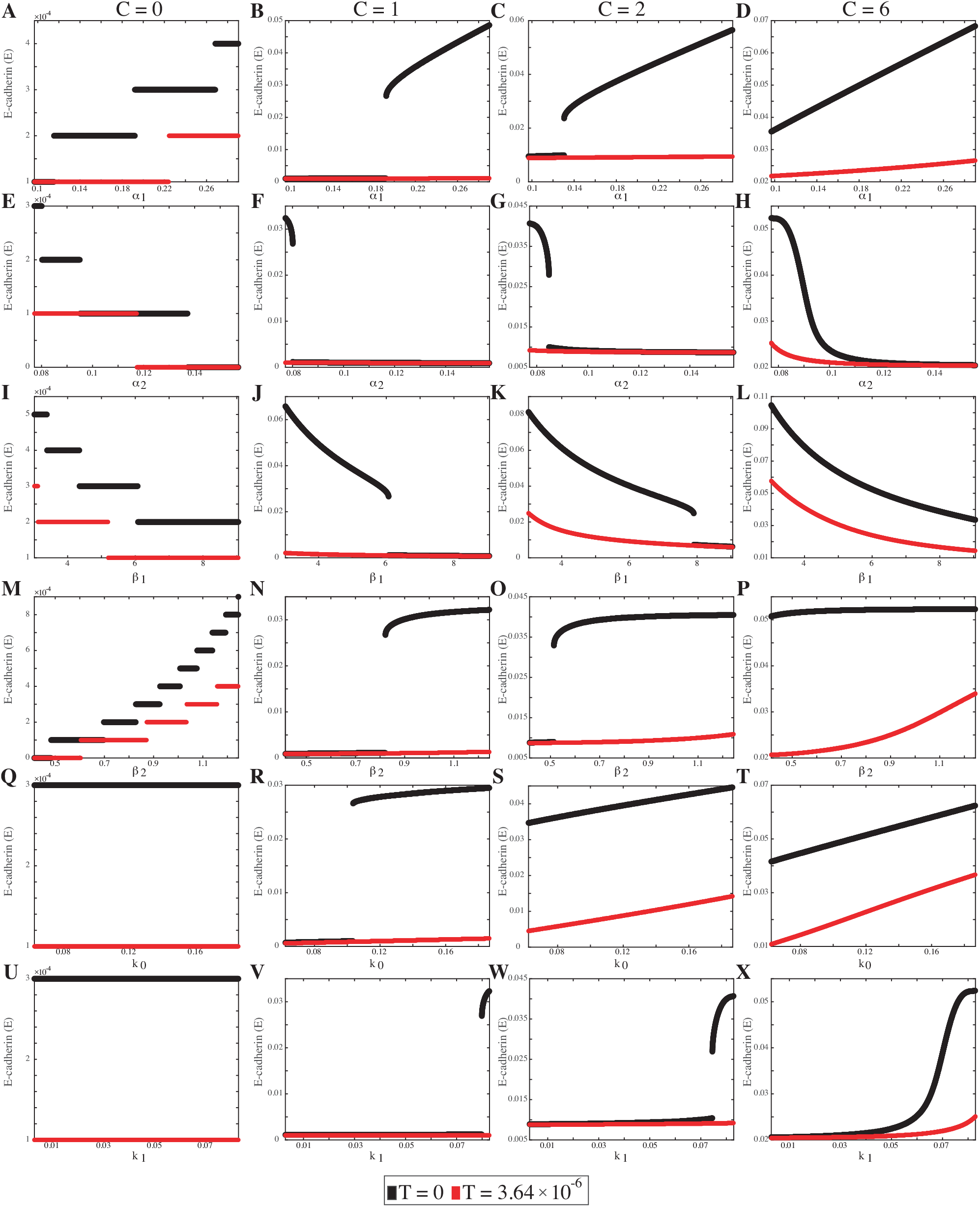
E-cadherin (E) monotonicity plots for parameters (A–D) *α*_1_, (E–H) *α*_2_, (I–L) *β*_1_, (M–P) *β*_2_, (Q–T) k0, and (U–X) *k*_1_ from Equations 1-2 with a ±50% parameter range. For each plot, 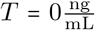 is in black and and 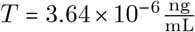 is in red. The level of cell–cell contact is given for each column with (A, E, I, M, Q, U) *C* = 0 cells, (B, F, J, N, R, V) *C* = 1 cells, (C, G, K, O, S, W) *C* = 2 cells, and (D, H, L, P, T, X) *C* = 6 cells.

**Figure S14:**
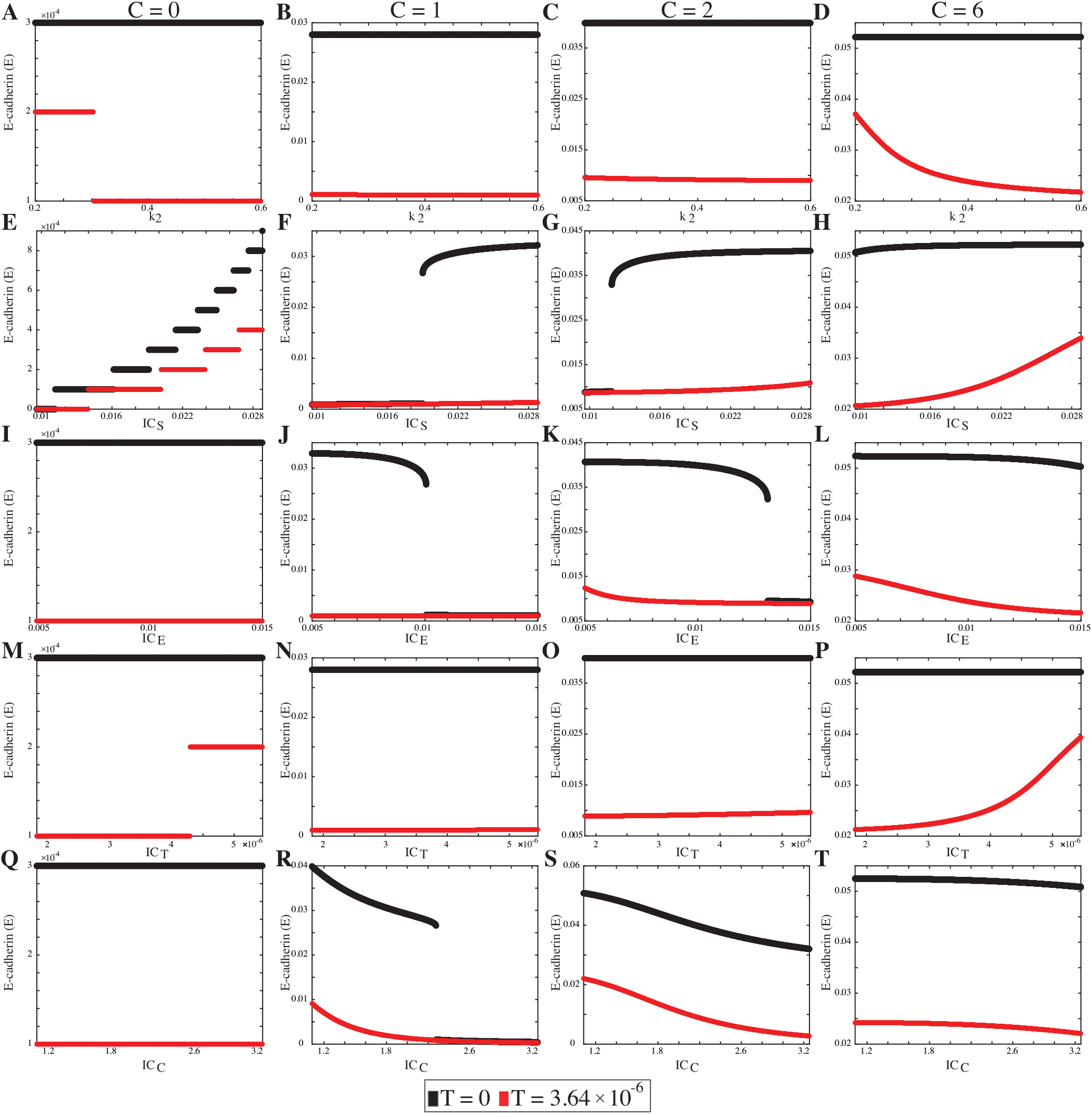
E-cadherin (E) monotonicity plots for parameters (A–D) k2, (E–H) ICS, (I–L) ICE, (M–P) ICT, and (Q–T) ICC from Equations 1-2 with a ±50% parameter range. For each plot, 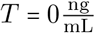 is in black and and 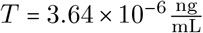 is in red. The level of cell–cell contact is given for each column with (A, E, I, M, Q) *C* = 0 cells, (B, F, J, N, R) *C* = 1 cells, (C, G, K, O, S) *C* = 2 cells, and (D, H, L, P, T) *C* = 6 cells.

**Figure S15:**
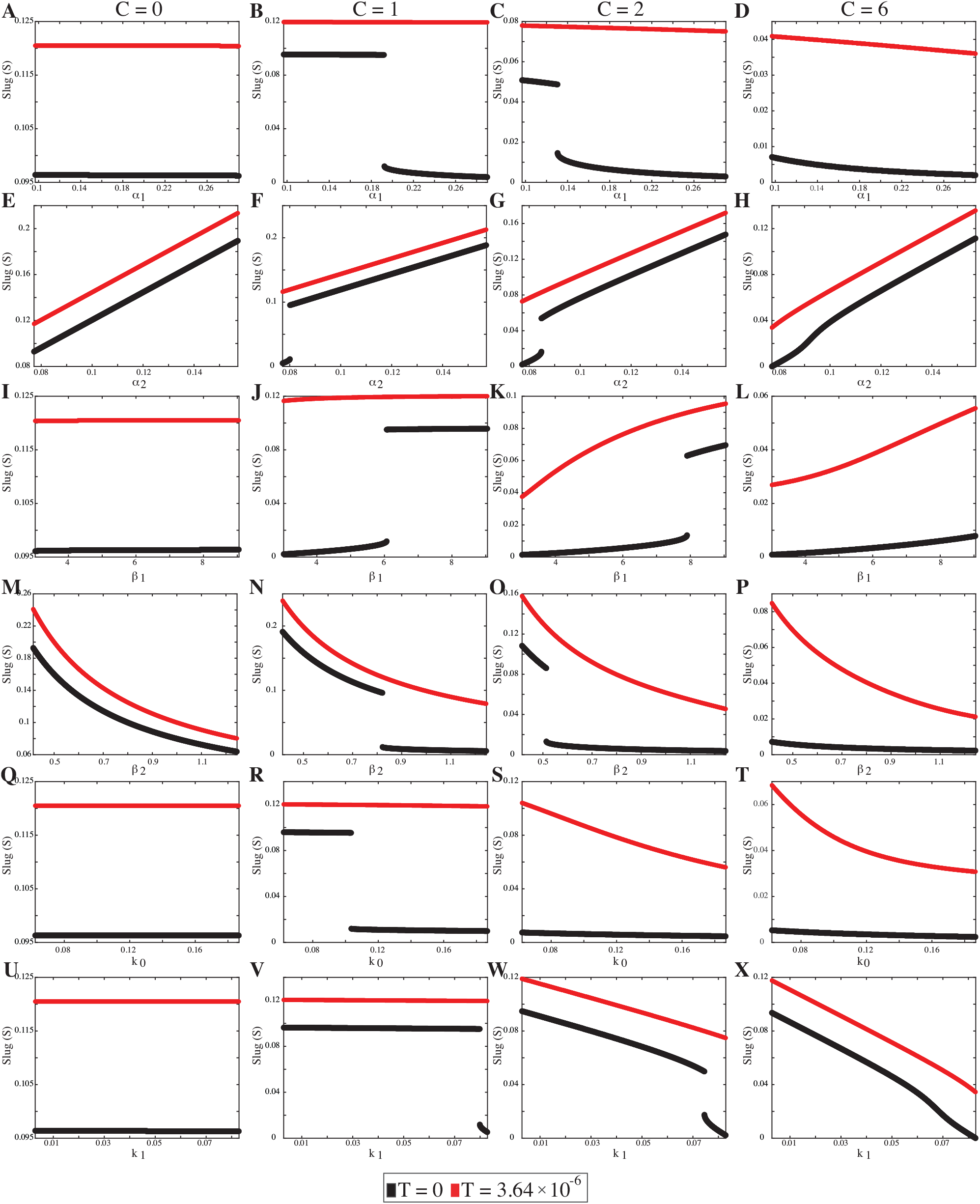
Slug (S) monotonicity plots for parameters (A–D) *α*_1_, (E–H) *α*_2_, (I–L) *β*_1_, (M–P) *β*_2_, (Q–T) k0, and (U–X) *k*_1_ from Equations 1-2 with a ±50% parameter range. For each plot, 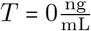 is in black and and 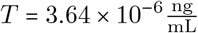 is in red. The level of cell–cell contact is given for each column with (A, E, I, M, Q, U) *C* = 0 cells, (B, F, J, N, R, V) *C* = 1 cells, (C, G, K, O, S, W) *C* = 2 cells, and (D, H, L, P, T, X) *C* = 6 cells.

**Figure S16:**
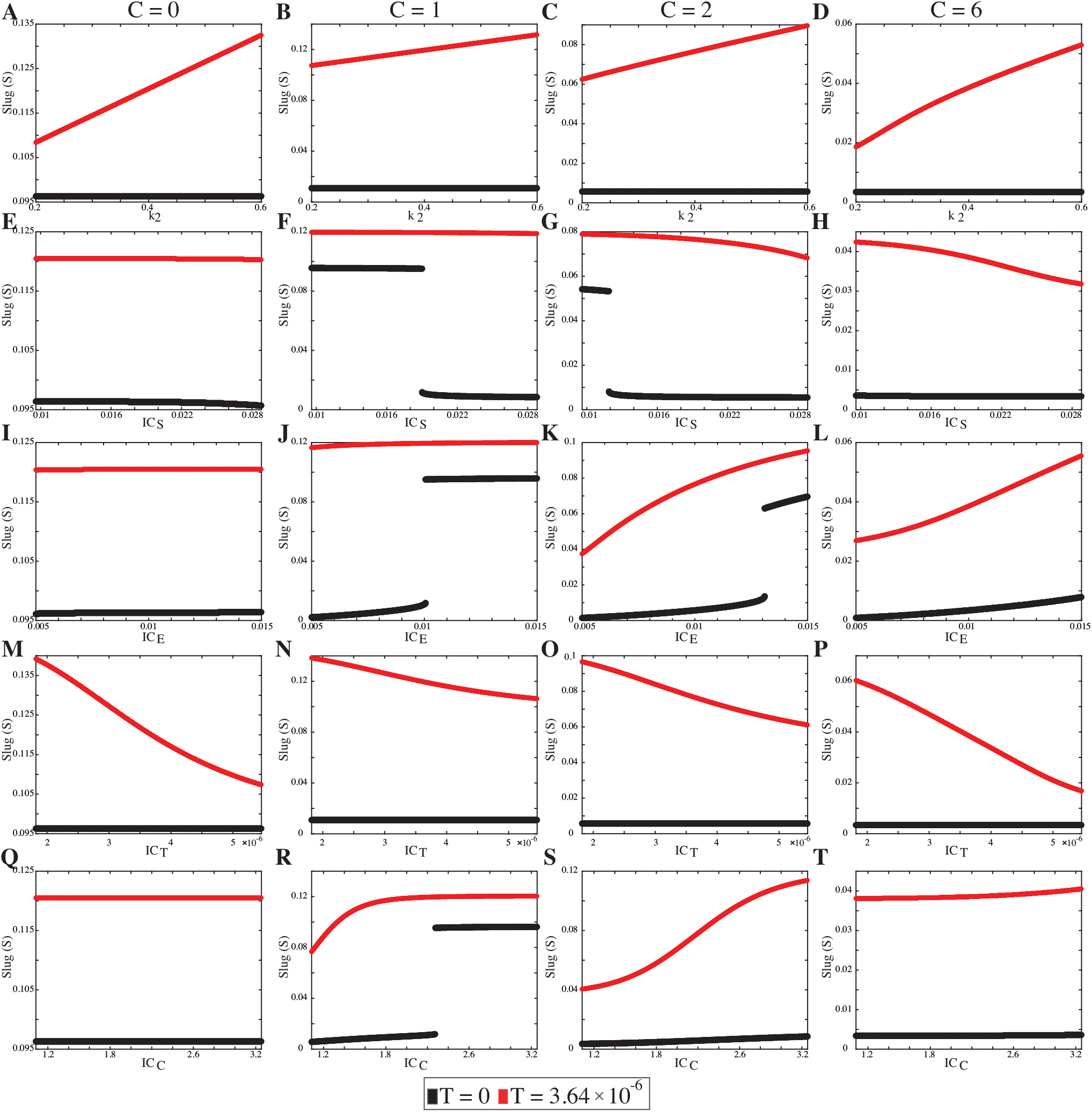
Slug (*S*) monotonicity plots for parameters (A–D) *k*_2_, (E–H) *IC*_*S*_, (I–L) *IC*_*E*_, (M–P) *IC*_*T*_, and (Q–T) *IC*_*C*_ from Equations 1-2 with a ±50% parameter range. For each plot, 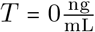 is in black and and 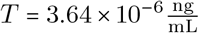 is in red. The level of cell–cell contact is given for each column with (A, E, I, M, Q) *C* = 0 cells, (B, F, J, N, R) *C* = 1 cells, (C, G, K, O, S) *C* = 2 cells, and (D, H, L, P, T) *C* = 6 cells.

**Table S1:** LHS-PRCC results for the ±10% parameter range as reported in [9].

| PRCC $\rho$ values for E-cadherin using $\pm 10\%$ Parameter Range | | | | | | | | |
| --- | --- | --- | --- | --- | --- | --- | --- | --- |
| Parameter | $T = 0 \frac{\text{ng}}{\text{mL}}$ | | | | $T = 3.64 \times 10^{-6} \frac{\text{ng}}{\text{mL}}$ | | | |
| | $C = 0$ | $C = 1$ | $C = 2$ | $C = 6$ | $C = 0$ | $C = 1$ | $C = 2$ | $C = 6$ |
| $\alpha_1$ | 0.1664 | 0.0650 | 0.0856 | 0.1058 | 0.1337 | 0.0240 | 0.0390 | 0.0468 |
| $\alpha_2$ | -0.4602 | -0.2684 | -0.4397 | -0.6594 | -0.2908 | -0.0564 | -0.0690 | -0.2615 |
| $\beta_1$ | -0.1827 | -0.2683 | -0.3052 | -0.2799 | -0.1311 | -0.2674 | -0.3597 | -0.6488 |
| $\beta_2$ | 0.5222 | 0.1451 | 0.1185 | 0.1179 | 0.3874 | 0.0789 | 0.0366 | 0.1329 |
| $k_0$ | -0.0085 | 0.1687 | 0.2170 | 0.1411 | -0.0157 | 0.2296 | 0.3439 | 0.5901 |
| $k_1$ | -0.0058 | 0.1168 | 0.3503 | 0.5946 | -0.0145 | 0.0052 | 0.0161 | 0.2228 |
| $k_2$ | 0.0050 | -0.0138 | 0.0038 | -0.0026 | -0.0610 | -0.0120 | -0.0196 | -0.0794 |
| $IC_S$ | 0.5038 | 0.1911 | 0.1176 | 0.1045 | 0.3963 | 0.0960 | 0.0297 | 0.1300 |
| $IC_E$ | -0.0062 | -0.0482 | -0.1188 | -0.0647 | -0.0038 | 0.0110 | -0.0160 | -0.0817 |
| $IC_T$ | -0.0007 | 0.0066 | -0.0050 | -0.0055 | 0.1113 | 0.0212 | 0.0196 | 0.1078 |
| $IC_C$ | 0.0098 | -0.7711 | -0.5167 | 0.0152 | 0.0160 | -0.9150 | -0.8481 | -0.0303 |
| PRCC $\rho$ values for Slug using $\pm 10\%$ Parameter Range | | | | | | | | |
| Parameter | $T = 0 \frac{\text{ng}}{\text{mL}}$ | | | | $T = 3.64 \times 10^{-6} \frac{\text{ng}}{\text{mL}}$ | | | |
| | $C = 0$ | $C = 1$ | $C = 2$ | $C = 6$ | $C = 0$ | $C = 1$ | $C = 2$ | $C = 6$ |
| $\alpha_1$ | 0.0056 | -0.0065 | -0.0458 | -0.0200 | 0.0033 | -0.0068 | -0.0263 | -0.0156 |
| $\alpha_2$ | 0.6684 | 0.7078 | 0.6281 | 0.7125 | 0.5643 | 0.5822 | 0.5149 | 0.6397 |
| $\beta_1$ | 0.0133 | 0.0318 | 0.2072 | 0.0887 | 0.0031 | 0.0072 | 0.2205 | 0.1822 |
| $\beta_2$ | -0.7206 | -0.6508 | -0.2685 | -0.1313 | -0.7708 | -0.7677 | -0.4358 | -0.3426 |
| $k_0$ | 0.0060 | 0.0035 | -0.1656 | -0.0360 | 0.0163 | 0.0020 | -0.2132 | -0.1551 |
| $k_1$ | -0.0012 | -0.0772 | -0.3898 | -0.6486 | 0.0107 | -0.0149 | -0.2093 | -0.5217 |
| $k_2$ | 0.0110 | -0.0001 | -0.0049 | 0.0020 | 0.1433 | 0.1312 | 0.1236 | 0.1543 |
| $IC_S$ | 0.0123 | -0.0622 | -0.0843 | -0.0268 | 0.0064 | -0.0026 | -0.0101 | -0.0179 |
| $IC_E$ | -0.0046 | 0.0278 | 0.2102 | 0.0744 | 0.0081 | 0.0064 | 0.2228 | 0.1458 |
| $IC_T$ | -0.0168 | 0.0012 | 0.0060 | 0.0052 | -0.2082 | -0.2026 | -0.1751 | -0.2381 |
| $IC_C$ | -0.0188 | 0.0505 | 0.3685 | -0.0224 | -0.0155 | 0.0263 | 0.5138 | 0.0002 |

## Notes

### Competing Interest Statement

The authors have declared no competing interest.

